# A novel framework to build saliva-based DNA methylation biomarkers: quantifying systemic chronic inflammation as a case study

**DOI:** 10.1101/2023.12.21.572866

**Authors:** Lisa J. Schmunk, Toby P. Call, Daniel L. McCartney, Hira Javaid, Waylon J. Hastings, Vanja Jovicevic, Dragoljub Kojadinović, Natacha Tomkinson, Eliska Zlamalova, Kirsty C. McGee, Jack Sullivan, Archie Campbell, Andrew M McIntosh, Veronika Óvári, Karl Wishart, Christian E. Behrens, Emma Stone, Miloš Gavrilov, Rob Thompson, Hurdle bio-infrastructure team, Thomas Jackson, Janet M. Lord, Thomas M. Stubbs, Riccardo E. Marioni, Daniel E. Martin-Herranz

## Abstract

Accessible and non-invasive biomarkers that measure human ageing processes and the risk of developing age-related disease are paramount in preventative healthcare. In this study, we describe a novel framework to train saliva-based DNA methylation (DNAm) biomarkers that are reproducible and biologically interpretable. By leveraging a reliability dataset with replicates across tissues, we demonstrate that it is possible to transfer knowledge from blood DNAm data to saliva DNAm data using DNAm proxies of blood proteins (EpiScores). We then apply these methods to create a new saliva-based epigenetic clock (InflammAge) that quantifies systemic chronic inflammation (SCI) in humans. Using a large blood DNAm human cohort with linked electronic health records and over 18,000 individuals (Generation Scotland), we demonstrate that InflammAge significantly associates with all-cause mortality, disease outcomes, lifestyle factors and immunosenescence; in many cases outperforming the widely used SCI biomarker C-reactive protein (CRP). We propose that our biomarker discovery framework and InflammAge will be useful to improve our understanding of the molecular mechanisms underpinning human ageing and to assess the impact of gero-protective interventions.

## 1. Introduction

DNA methylation (DNAm) has emerged as a powerful data type to build biomarkers that quantify the risk for developing age-related diseases and the ageing process itself (Horvath & Raj, 2018). This may be due to the ability of DNAm to capture information from both genetics and lifestyle/environmental factors (van Dongen et al., 2016). DNAm patterns associated with ageing and maximum lifespan are evolutionarily conserved (Haghani et al., 2023; Lu et al., 2023), suggesting a critical role in the biology of ageing.

Epigenetic ageing clocks are becoming more widespread in clinical research, including in the context of quantifying the effect of candidate healthspan extending interventions in human trials (Moqri et al., 2023), and in the consumer market for preventative healthcare applications (Knoppers et al., 2021). This has been driven by significant improvements in the design and performance of these biomarkers, both from a technical and a clinical perspective. Early examples include the multi-tissue Horvath clock (Horvath, 2013), the blood-specific Hannum clock (Hannum et al., 2013) or the skin-blood clock (which has good predictive performance in saliva samples and *in vitro* cell culture) (Horvath et al., 2018). These were trained directly on chronological age and are able to predict it with remarkable accuracy, which has applications, for example, in the forensic field (Ambroa-Conde et al., 2022).

Second-generation epigenetic clocks have improved associations with clinical outcomes (Margiotti et al., 2023). Instead of training DNAm biomarkers directly on chronological age, it is possible to train them on intermediate phenotypic proxies that better capture morbidity and all-cause mortality risk. These include phenotypic ageing measurements derived from clinical biomarkers (PhenoAge) (Levine et al., 2018), a proxy that quantifies the pace of ageing across multiple organs/systems (DunedinPACE) (Belsky et al., 2022) or DNAm-based predictors of plasma proteins and smoking exposure that accurately predict all-cause mortality (GrimAge) (Lu et al., 2019). Further methodological innovations, such as those that leverage principal component analysis (PCA) to reduce technical variability of CpG probes, have enhanced the test-retest reliability of DNAm biomarkers, which is critical to capture longitudinal trajectories and intervention effects (Higgins-Chen et al., 2022).

Despite these improvements, two problems still remain for existing DNAm-based biomarkers: (1) the lack of biological interpretability with current epigenetic clocks behaving like a ‘black box’ (with the exception of perhaps GrimAge, where features can be interpreted), and (2) the paucity of second-generation reproducible epigenetic clocks in saliva (which is a more accessible and scalable sample type for preventative healthcare propositions when compared to blood). In this work, we outline a novel framework to build saliva-based DNAm biomarkers that addresses both of these challenges by leveraging a library of blood-based DNAm predictors of different plasma proteins (protein epigenetic scores or EpiScores) (Aslibekyan et al., 2018; Gadd et al., 2022; Hillary et al., 2020; Ligthart et al., 2016; Stevenson et al., 2020, 2021).

Ageing is driven by many molecular processes known as ‘Hallmarks of Ageing’ (López-Otín et al., 2023), with the geroscience hypothesis postulating that measuring and then targeting these processes rather than individual diseases will lead to a more successful impact on human healthspan (Kennedy et al., 2014; Sierra, 2016). Low-grade systemic chronic inflammation (SCI), also known as ‘inflammageing’, has recently emerged as one of these hallmarks (López-Otín et al., 2023). SCI is characterised by persistent and non-resolving inflammation in the absence of an acute trigger, leading to collateral tissue damage and an increase in disease risk (Furman et al., 2019).

A subset of blood-based biomarkers traditionally associated with acute inflammatory responses have been reported to also consistently change their baseline levels during human ageing; thus they have been proposed as SCI biomarkers. This includes biomarkers such as C-reactive protein (CRP) (Tang et al., 2018), erythrocyte sedimentation rate (ESR) (Siemons et al., 2014), IL-6 (Stevenson et al., 2021), TNF-α (Koelman et al., 2019), interferon-gamma (IFN-γ) (Koelman et al., 2019), CXCL9 (Sayed et al., 2021) and other cytokines and chemokines (Koelman et al., 2019). However, due to the high biological variability of these biomarkers with acute inflammation and infection status, single measurements are normally insufficient to determine the SCI status of an individual.

Recently, omics-based approaches have used machine learning algorithms to predict SCI status. Notable examples include leveraging the immunoglobulin G (IgG) glycome (Štambuk et al., 2020) or iAge (which integrates different inflammatory markers into a single metric) (Sayed et al., 2021). However, these biomarkers still require invasive procedures to access blood, they may not be very stable (e.g. cytokines in iAge) and they require access to expensive and complex molecular assays (which are not globally available across laboratories); all of which increase the barriers for widespread adoption in preventative healthcare settings.

In this work, we demonstrate the utility of our framework to train a new saliva-based DNAm biomarker for SCI (which we have named InflammAge). InflammAge addresses the hurdles outlined above for SCI quantification while demonstrating high technical reliability, significant associations with clinical outcomes and increasing biological interpretability when compared to other epigenetic clocks.

## 2. Results

### A novel framework to train biologically-interpretable and reproducible DNAm biomarkers in saliva

To harness the power of DNA methylation (DNAm) as a data type and saliva as an easily accessible human sample, we created a framework for developing novel saliva-based DNAm biomarkers informed by biological interpretability.

Genome-wide DNAm can be quantified using Illumina Infinium methylation microarrays. The most widely used technology is the Illumina Infinium MethylationEPIC v1.0 BeadChip array (EPIC), which quantifies ∼850,000 CpG sites. Although the EPIC array is designed to enhance novel discovery for a broad consumer base, we aimed to create a more cost-effective approach to DNAm assessment at sites relevant to geroscience. To generate this custom array, termed the Hurdle DNAm platform, we first selected a set of CpG sites with strong phenotypic associations established by published epigenome-wide association studies (EWAS) (Battram et al., 2022; Li et al., 2019). We enriched for actionable traits, lifestyle and environmental exposures, epigenetic clocks and complex diseases; resulting in > 30,000 CpG sites that are good candidate features for biomarker development (see Methods and **Supplementary Fig. S1**).

EpiScores are DNAm-based predictors of varied traits with direct biological interpretability, such as blood protein levels. They have been created based on blood DNAm EWAS of large cohorts with matched data types (Gadd et al., 2022; Stevenson et al., 2020, 2021). As DNAm captures cellular states upstream of gene regulation, these EpiScores may act as de-noised proxies for the blood proteins, better capturing chronic disease risk. This may prevent confounding with short-term fluctuations in blood concentrations, for example, due to circadian rhythms (Nilsonne et al., 2016) or acute infection (Largman-Chalamish et al., 2022). We propose that these EpiScores can be transferred from blood (where they were originally trained) to saliva (given the tissue similarity, see Discussion); and then used as improved biological features to train machine learning models that are able to predict complex phenotypes from DNAm data (while still allowing for biological interpretability).

Our DNAm biomarker discovery framework consists of the following steps (**Fig 1a**.):

1. generate genome-wide DNA methylation data in all datasets;
2. calculate a library of published blood EpiScores as features in all datasets;
3. filter the calculated features in the training dataset by reproducibility across tissues using a reliability dataset with matched blood and saliva DNAm data;
4. filter the remaining features in the training dataset by technical reliability of each feature within the tissue both within blood and within saliva, using technical replicates from a reliability dataset;
5. select final features in the training dataset based on biological criteria (e.g. association with physiological ageing, association with a disease process, etc);
6. train the final machine learning model on the desired phenotypic outcome (e.g. chronological age, all-cause mortality, time-to-disease, etc) using the selected features in the training dataset in order to generate the DNAm biomarker; and
7. validate the final DNAm biomarker in independent test datasets.

**Fig. 1.**
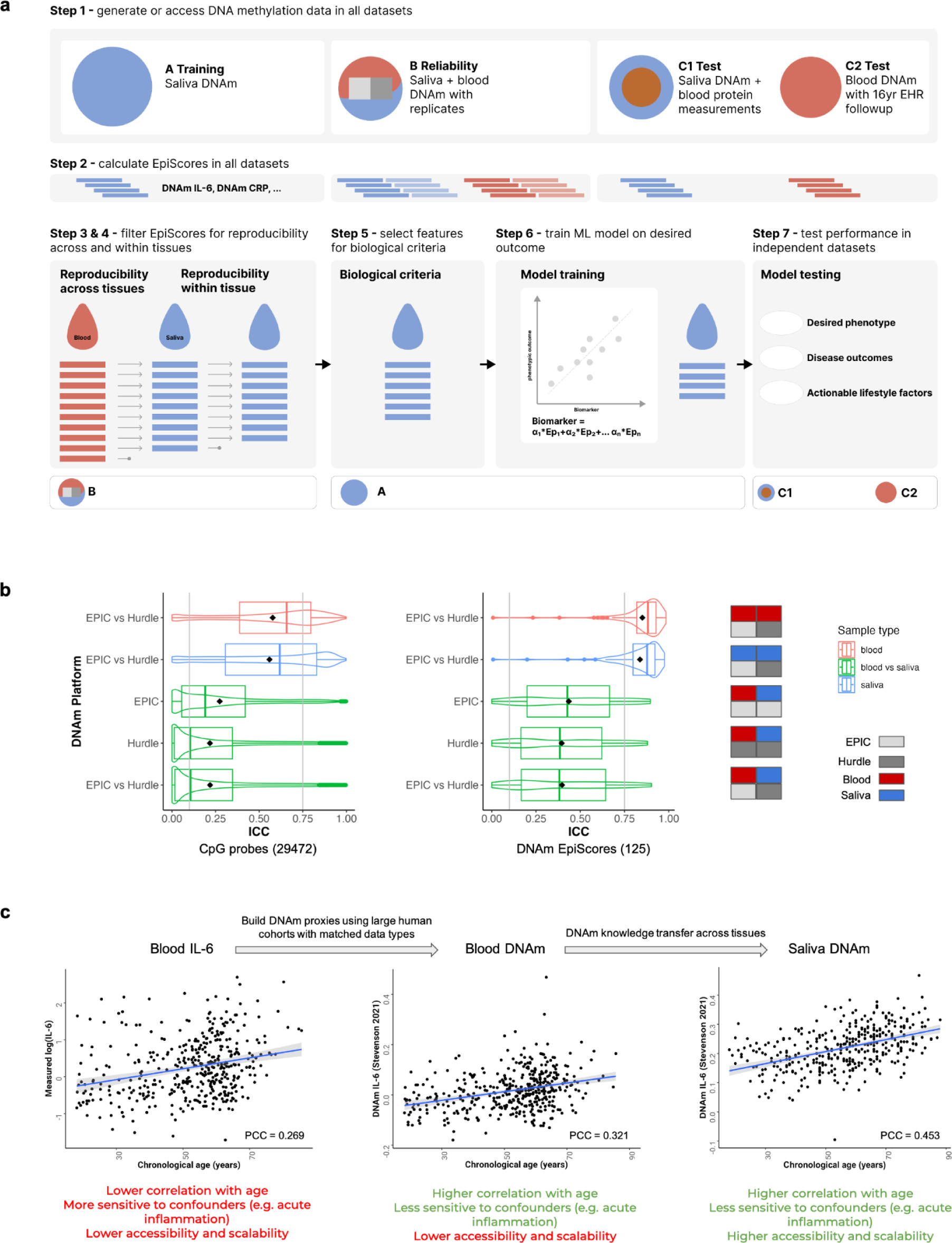
A novel framework to train biologically-interpretable and reproducible DNAm biomarkers in saliva. a. Schematic showing the steps (1-7) and datasets (training, reliability, test) required in the framework to train novel DNAm biomarkers in saliva. EHR = electronic health records, IL-6 = Interleukin 6, CRP = C-reactive protein. **b** Reproducibility analysis for the CpG probes (left) and EpiScore (right) features considered during the training of InflammAge. The distributions of intraclass correlation coefficients (ICC) were calculated in the reliability dataset and displayed as box and violin plots for five different comparisons (including within and between blood and saliva tissues). The box plots show the interquartile range (IQR) as squares and the mean (black diamond). Grey vertical lines indicate ICC filtering thresholds at 0.75 for within tissues and 0.10 for between tissues. **c** Plots showing the transfer of biological information from blood proteins (left, data from (Stevenson et al., 2021)), to blood DNA methylation EpiScores (center, data from (Stevenson et al., 2021)) and finally to saliva DNA methylation EpiScores (right, data from the saliva training dataset). In the case of InflammAge, saliva DNAm SCI features were selected based on the association with ageing (chronological age). The SCI biomarker IL-6 is used as an example, which shows consistently higher levels with increased age.

In the next section, we describe how this hypothesis-driven framework can be applied to build a novel saliva-based DNAm biomarker of systemic chronic inflammation (SCI) in humans (InflammAge).

### Applying the framework to train a saliva-based DNAm biomarker for systemic chronic inflammation (InflammAge)

Different proteins (including cytokines) have been associated with SCI in the literature, providing evidence that this ageing hallmark can be quantified at the molecular level in blood. Furthermore, we have also observed the age-related association of many of these biomarkers in one of our healthy ageing cohorts (see Alpha test dataset in Methods), which included an increase in the baseline levels of GDF-15 (Spearman correlation coefficient SCC=0.76, FDR-adj. p-value=3.8x10^-12^), CXCL9 (SCC=0.67, FDR-adj. p-value=6.6x10^-9^), TNF-α (SCC=0.40, FDR-adj. p-value=4.9x10^-3^), IL-6 (SCC=0.36, FDR-adj. p-value=0.013), CRP (SCC=0.33, FDR-adj. p-value=0.027) or CCL11 (SCC=0.30, FDR-adj. p-value=0.0498) during ageing (**Supplementary Fig. S2**). Based on this, we collated a library of published blood DNAm CpG sites and EpiScores that are associated with cytokines and other inflammation-related proteins. This includes CRP (Ligthart et al., 2016; Stevenson et al., 2020), IL-6 (Stevenson et al., 2021), TNF-α (Aslibekyan et al., 2018), CXCL9, CCL11, IL-18R1 (Hillary et al., 2020) and 109 EpiScores containing inflammatory proteins measured in the Olink and SOMAscan platforms (Gadd et al., 2022). These features were calculated across all DNAm samples in all datasets (training, reliability, test; step 2 of the framework).

To further ensure the reproducibility of the features going into the training stage, we generated a reliability dataset with DNAm technical replicates for matched saliva and blood samples by measuring methylation profiles twice on the Illumina EPIC array and our custom Hurdle DNAm platform (see Methods). We calculated intraclass correlation coefficients (ICCs) between individual CpG probes present on both platforms (29,472) and for each summary EpiScore (125 in total) for the following comparisons: blood EPIC vs blood Hurdle, saliva EPIC vs saliva Hurdle, blood EPIC vs saliva EPIC, blood Hurdle vs saliva Hurdle and blood EPIC vs saliva Hurdle (**Fig. 1b**). Using this approach, we were able to validate which CpG probes and EpiScores could be measured reliably across platforms and tissues. We observed that EpiScores showed higher reliability both within tissues (with mean values of ICC_Blood_=0.85, 95% CI [0.76-0.90]; ICC_Saliva_=0.84, 95% CI [0.74-0.90]) and between tissues (mean ICC_Blood-vs-Saliva_=0.41, 95% CI [0.13-0.60]) compared to means of individual CpG probes (ICC_Blood_=0.58, 95% CI [0.42-0.69]; ICC_Saliva_=0.56, 95% CI [0.40-0.68]; ICC_Blood-vs-Saliva_=0.23, 95% CI [0.02-0.40]). Furthermore, many EpiScores showed strong positive correlations between blood EPIC and saliva Hurdle replicates collected from the same individual at the same time (**Supplementary Fig. S3a**), adding robust evidence to the ability to transfer EpiScores information from blood DNAm to saliva DNAm data. We therefore decided to train our InflammAge clock using the EpiScores instead of the individual CpGs that make up the EpiScores.

Using the V1 samples from the reliability dataset (see Methods), different ICC thresholds for the EpiScores were benchmarked in order to optimise the ICC for the final InflammAge biomarker (**Supplementary Fig. S4**). This resulted in selecting EpiScores with ICC >0.10 between blood and saliva (step 3 of the framework; 101/125 EpiScores remained) and an ICC >0.75 within saliva and within blood replicates (step 4 of the framework; 86/125 EpiScores remained). The 86 EpiScores with high reliability were used downstream for training.

To train our InflammAge biomarker in saliva, we assembled a cohort with human saliva DNAm samples from published and in-house data (training dataset, see Methods). This dataset (N=338, 27% female) covers the adult human lifespan (age range of 18-88 years, median 58.5 years, mean = 56.8 years), including individuals from diverse genetic backgrounds (see Methods, **Supplementary Table S1**). In this cohort, we calculated the correlation (Pearson correlation coefficient, PCC) of each EpiScore with chronological age in saliva (**Fig. 2a**). The biological assumption is that if the EpiScore for an inflammatory protein increases or decreases in the human population (i.e. cross-sectionally) with age, then it is more likely to be a marker for SCI. We kept those features with associations with age in saliva that had an FDR-adj. p-value <0.05 (step 5 of the framework, see **Fig. 1c**).

**Fig. 2.**
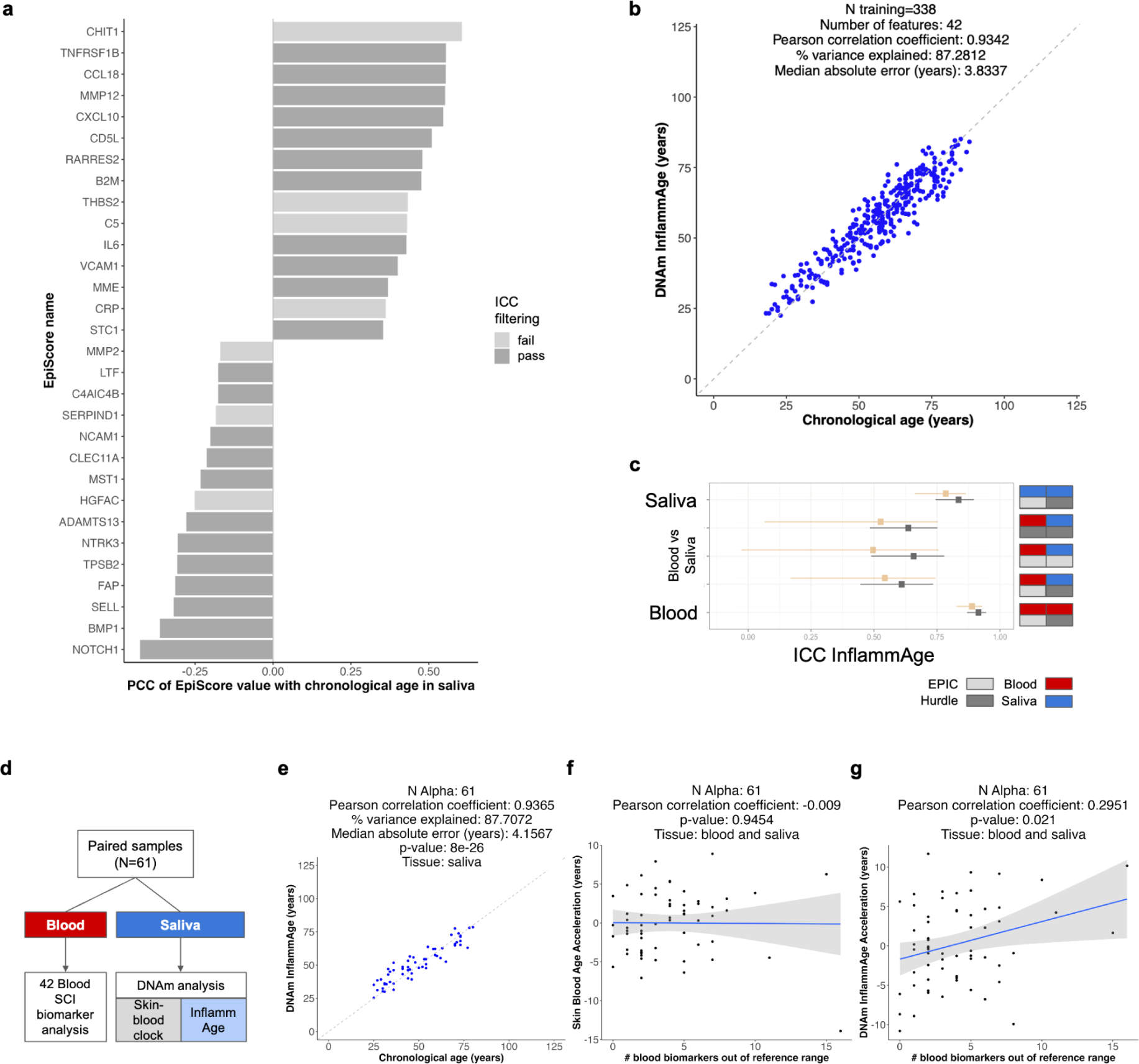
Train and test results of a saliva-based DNAm biomarker for systemic chronic inflammation (InflammAge). a. Barplot displaying the top 15 positive and top 15 negative SCI DNAm EpiScore features that are significantly associated with chronological age (FDR-adj. p-value<0.05) in the training dataset in saliva. Features that passed all technical filtering steps (dark-grey bars) were used to train InflammAge. PCC: Pearson correlation coefficient. **b** Scatterplot showing the performance of InflammAge in the training dataset. **c** Reproducibility of InflammAge within and between blood and saliva tissues as measured by intraclass correlation coefficient (ICC) in the reliability dataset (using V2 samples). The square represents the mean ICC value and the error bars the 95% confidence intervals. ICC values in black show the improvement achieved by filtering features using ICCs before training (steps 3 and 4 of the framework). Results for an alternative InflammAge model without ICC-based filtering are shown in orange. The legend on the right represents the type of reliability comparison: the upper row represents the tissues (red: blood, blue: saliva) and the lower row the DNAm platform (light-grey: EPIC, dark-grey: Hurdle). **d-g** Testing InflammAge performance in an independent test cohort (Alpha). **d** Alpha cohort design (N=61 individuals) with matched saliva DNAm and gold standard blood SCI biomarkers. **e** Scatterplot showing the performance of InflammAge in the Alpha test dataset. **f-g** Scatterplots showing the association between SCI status in blood (quantified as the total number of SCI blood biomarkers outside of the normal reference ranges in an individual) with f) age acceleration in the saliva-trained skin-blood epigenetic clock (Horvath et al., 2018) and g) age-adjusted InflammAge acceleration. Blue lines indicate a linear model fit and grey shading the confidence interval.

After this filtering step, we retained 63/125 EpiScores; with a total of 42 showing a positive correlation with age and a total of 21 negatively correlated with age. The top 15 EpiScore correlation coefficients are displayed in **Fig. 2a**. EpiScores with positive correlation with age in saliva included well characterised pro-inflammatory markers like TNFRSF1B (PCC=0.56, FDR-adj. p-value=4.8x10^-27^), CCL18 (PCC=0.56, FDR-adj. p-value=4.8x10^-27^), MMP12 (PCC=0.55, FDR-adj. p-value=6.8x10^-27^), CXCL10 (PCC=0.55, FDR-adj. p-value=2.8x10^-26^), or IL-6 (PCC=0.43, FDR-adj. p-value=1.7x10^-15^) (see Discussion). EpiScores with negative correlation with age in saliva included NOTCH1 (PCC=-0.43, FDR-adj. p-value=2.1x10^-15^), BMP1 (PCC=-0.36, FDR-adj. p-value=4.3x10^-11^), FAP (PCC=-0.31, FDR-adj. p-value=1.9x10^-8^) and NTRK3 (PCC=-0.31, FDR-adj. p-value=3.9x10^-8^). On conducting a gene ontology analysis, we found that positively age-associated EpiScore genes were significantly enriched for inflammatory processes, cytokine activity and response to stress (**Supplementary Fig. S5a**). Negatively age-associated EpiScore genes were significantly associated with serine peptidase activity, humoral immune and biotic stimulus response (**Supplementary Fig. S5b**). These results give further confidence that the proteins quantified by the selected EpiScores are indeed enriched for inflammatory and immune response pathways.

Next, we trained our DNA methylation biomarker (InflammAge) in our saliva training dataset using an elastic net algorithm (step 6 of the framework, see Methods). We selected the 63 DNAm features enriched for inflammation-related pathways as the input and chronological age as the outcome. In the training dataset, InflammAge predicted chronological age with a performance in range with gold standard epigenetic clocks (PCC=0.9342, median absolute error (MAE)=3.83 years, **Fig. 2b**). A total of 42 DNAm features (containing a total of 3788 unique CpG sites in the Hurdle DNAm platform) constitute the final InflammAge model.

We then tested the performance of InflammAge in two independent datasets (step 7 of the framework). In the reliability dataset (V2 samples), performance of InflammAge was good in saliva (ICC=0.84, 95% CI [0.74-0.90]) and excellent in blood (ICC=0.91, 95% CI [0.87-0.95]) (**Fig. 2c**), showing that saliva DNAm biomarkers are reproducible when trained using our framework. Moderate reliability (ICC=0.61, 95% CI [0.44-0.73]; for saliva Hurdle vs blood EPIC) between tissues was also observed, which highlights that even though InflammAge was trained in saliva DNAm samples, the results can be extrapolated to blood DNAm samples assessed on alternative platforms (**Supplementary Fig. S3** and **Supplementary Fig. S4**). Importantly, training an alternative InflammAge model using all the features associated with age as input but without filtering for ICC (i.e., skipping steps 3 and 4 in the framework) resulted in a less reliable biomarker with wide confidence intervals across all comparisons (**Fig. 2c**). This validates the importance of the reliability dataset and steps 3 and 4 in our framework.

Finally, we tested the performance of InflammAge in the Alpha test dataset (see Methods). This cohort was independently recruited for this work (N=61, 46% female) and also covers the adult human lifespan (age range of 25-80 years, median = 46 years, mean = 49.57 years), including only self-reported healthy individuals (see Methods for full exclusion criteria). Therefore, this dataset should give a fair assessment of the performance of InflammAge during human physiological ageing. Importantly, this cohort included matched saliva DNAm data and blood SCI biomarkers (such as cytokines and other inflammatory proteins; **Fig. 2d**), which allows for assessing whether InflammAge is capturing changes in the SCI status of the individuals (besides tracking chronological age). InflammAge predicted chronological age with a correlation (PCC) of 0.94 and MAE of 4.2 years (**Fig. 2e**). Moreover, we calculated age-adjusted InflammAge acceleration (see Methods), which represents the difference between measured InflammAge and the average InflammAge expected for someone with the chronological age of the measured individual. A positive (>0) InflammAge acceleration should be associated with higher SCI when compared to other individuals of the population with the same chronological age (i.e., that person should be ‘inflammageing’ faster). Indeed, InflammAge acceleration was associated with the inflammatory biomarker CRP (measured by ELISA from blood serum, PCC=0.28, p-value=0.03, **Supplementary Fig. S6**) and also associated with the total number of blood SCI biomarkers outside of reference ranges (PCC=0.30, p-value=0.02, **Fig. 2g**). On the contrary, another saliva-based epigenetic clock (skin-blood clock) (Horvath et al., 2018) was not correlated (PCC=-0.01, p-value=0.95, **Fig. 2f**). This demonstrates the ability of InflammAge to capture SCI status from a saliva DNAm sample.

### InflammAge is accelerated in SCI-related clinical outcomes

Since saliva DNAm cohorts with mapped clinical outcomes are rare, we tested InflammAge in Generation Scotland (GS), one of the world’s largest biobanks with access to blood DNAm samples matched to electronic health records (see Methods). GS has blood DNAm samples from 18,865 individuals (58.8% female) covering the adult human lifespan (age range of 18-99 years, median 49.2 years, mean=47.8 years). More summary information on the GS cohort is provided in **Supplementary Table S2**.

First, we calculated InflammAge in blood DNAm samples for 18,865 individuals in the GS cohort. InflammAge showed a strong correlation with chronological age in the GS dataset, although an intercept shift was observed (i.e. a systematic underestimation of InflammAge in blood when compared to saliva DNAm samples; PCC=0.904, MAE=10.42 years; **Supplementary Fig. S7**). Given these results, we decided to proceed to use GS as a test dataset using age-adjusted InflammAge acceleration, which would correct for such a shift in the downstream analyses (see Methods).

We tested the Spearman correlation coefficients (SCC) between InflammAge acceleration and basic demographic and biochemistry variables (**Supplementary Table S3**). InflammAge acceleration showed a significant difference when stratified by sex, where males had a mean InflammAge acceleration of 0.198 compared to -0.139 years in females (t-test, p-value=3.1x10^-4^). This is consistent with reports in the literature that highlight higher SCI baseline levels in males (Martínez de Toda et al., 2023). InflammAge acceleration (from samples taken between 2006 and 2011) was also associated with positive COVID-19 disease diagnosis, ascertained from primary and secondary care records (HR=1.24, 95% CI [1.1-1.4], p-value=3.1x10^-4^).

Next, we screened for an association between age-adjusted InflammAge acceleration and 308 diseases that were annotated using disease codes from Kuan et al. (Kuan et al., 2023). This was performed using Cox proportional hazards regression analyses that included demographic risk factors (age, sex, BMI, years of education, socioeconomic status, smoking pack-years, and DNAm processing batch) as covariates. This allows us to assess if InflammAge adds significant value when modelling the risk of disease when compared with traditional risk factors. InflammAge acceleration was associated with the incidence of 18 diseases after multiple testing correction and without violation of the model assumptions (**Fig. 3a)**; 33 diseases were significant when using unadjusted p-values (**Supplementary Table S4**). The 18 hits included strong associations with the risk of developing diseases where SCI plays an important role in the aetiology, such as cancer (e.g. non-Hodgkin’s lymphoma, HR=1.47, 95% CI [1.20-1.81], FDR-adj. p-value=5.78x10^-04^) (Makgoeng et al., 2018), diseases of the circulatory system (e.g. peripheral arterial disease, HR=1.24, 95% CI [1.09-1.41], FDR-adj. p-value=3.37x10^-3^) (Brevetti et al., 2010), diseases of the digestive system (e.g. cholecystitis, HR=1.25, 95% CI [1.13-1.40], FDR-adj. p-value=1.01x10^-04^) (Jones et al., 2023) and diseases of the respiratory system (e.g. COPD, HR=1.23, 95% CI [1.13-1.35], FDR-adj. p-value=2.06x10^-5^) (Oudijk et al., 2003). InflammAge acceleration outperformed blood CRP (a gold standard inflammatory biomarker) in 15 out of these 18 diseases (based on a comparison of FDR-adj. p-values). Furthermore, InflammAge acceleration outperformed the only widely available saliva-based ageing DNA methylation biomarker (skin-blood epigenetic clock) (Horvath et al., 2018) in all these 18 diseases. CRP generally outperformed InflammAge in the “diseases of the circulatory system” category and skin-blood was mainly predictive for some cancer types. This confirms that InflammAge captures SCI-related disease outcomes from an accessible tissue.

**Fig. 3.**
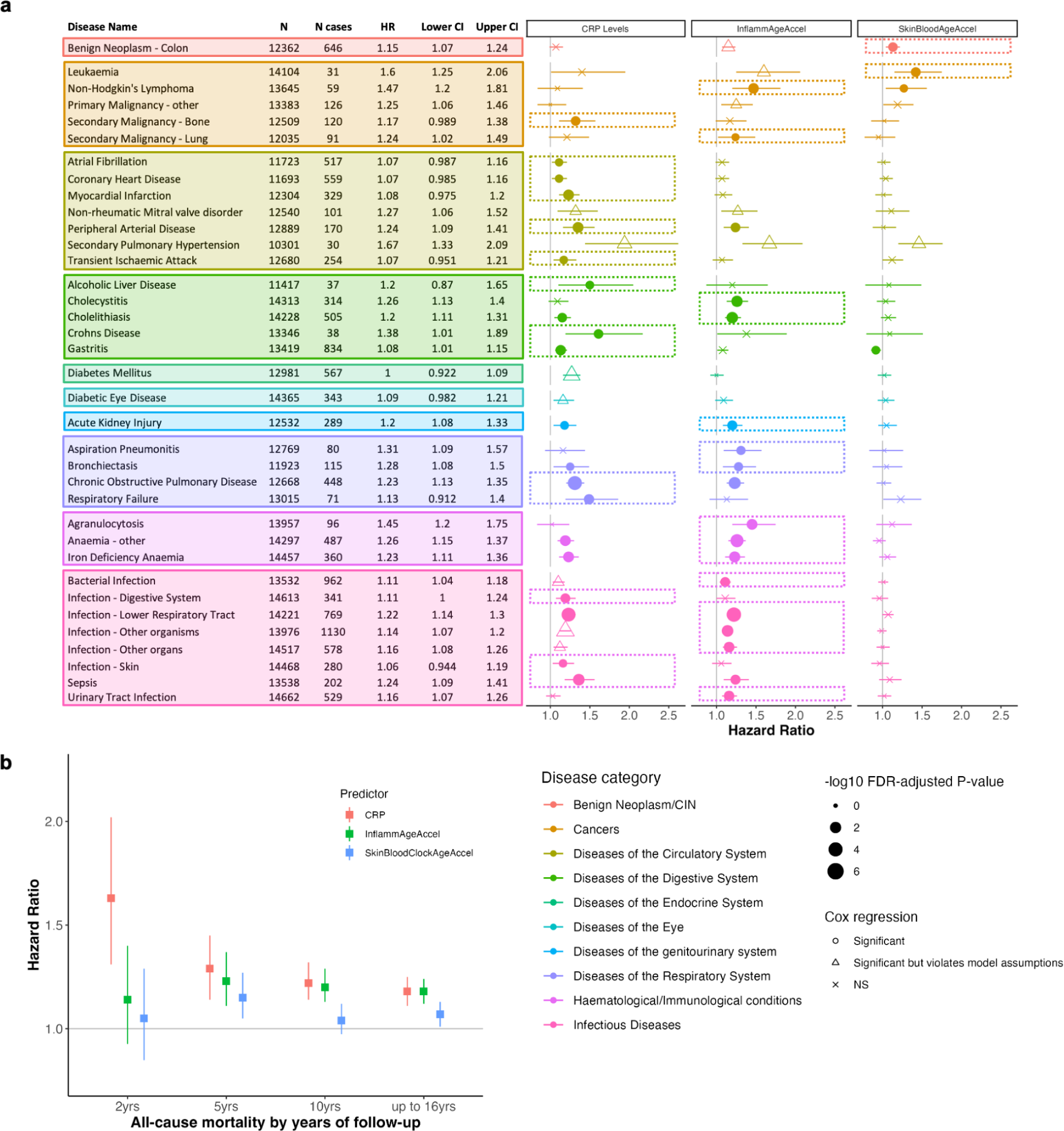
InflammAge acceleration performance in clinical endpoints when compared to gold standard SCI markers in Generation Scotland. Incidence of clinical endpoint was predicted for fully adjusted InflammAge acceleration (InflammAgeAccel, see Methods), skin-blood clock age acceleration (SkinBloodClockAgeAccel) (Horvath, 2013) and levels of a gold standard SCI biomarker (CRP, as measured by ELISA in GS). **a** Results of a disease incidence Cox regression analysis (see Methods). Displayed are 36 diseases with >30 cases in GS and association FDR-adj. p-value<0.05 for at least one of the three predictors. The table shows the hazard ratio (HR) and 95% confidence intervals per disease for InflammAgeAccel. Full summary statistics are available in **Supplementary Fig. S4**. The plot shows hazard ratios and 95% confidence intervals across all three predictors with the most significant one per disease highlighted by dashed boxes. The shapes of the data points indicate the significance and violations of model assumptions (see Methods); NS = non-significant. The sizes of the data points reflect the level of significance (proportional to -log10 of FDR-adj. p-value) with larger symbols having lower p-values. Data points are coloured by disease category from Kuan et al. (Kuan et al., 2023). Filled symbols indicate significance without model assumption violations. **b** Mortality prediction in the Generation Scotland cohort for 2 years (N=66 events), 5 years (N=241 events), 10 years (N=683 events) and the entire follow-up period (N=1,031 events). Squares show the hazard ratio and lines show 95% confidence intervals.

Cox proportional hazards regression models were performed to assess the relationship between InflammAge acceleration (and other available biomarkers, such as blood CRP and skin-blood epigenetic clock) and all-cause mortality. This was assessed at intervals of 2 years (N=66 events), 5 years (N=241 events), 10 years (N=683 events), and the entire follow-up period (up to 16 years, N=1031 events). Except for the 2-year interval, there was a significant relationship between InflammAge acceleration and all-cause mortality (**Fig. 3b**). The performance of InflammAge acceleration was comparable to that of blood-based CRP measurements and outperformed the skin-blood clock when predicting mortality within 5 years (HR_CRP_=1.29, 95% CI [1.14-1.45], HR_InflammAgeAccel_=1.23, 95% CI [1.11-1.37], HR_SkinBloodAgeAccel_=1.15, 95% CI [1.05-1.27]), 10 years (HR_CRP_=1.22, 95% CI [1.14-1.32], HR_InflammAgeAccel_=1.20, 95% CI [1.13-1.29], HR_SkinBloodAgeAccel_=1.04, 95% CI [0.97-1.12]) and the full follow-up period (up to 16 years: HR_CRP_=1.18, 95% CI [1.11-1.25], HR_InflammAgeAccel_=1.18, 95% CI [1.12-1.24], HR_SkinBloodAgeAccel_=1.07, 95% CI [1.01-1.13]).

### InflammAge acceleration is associated with lifestyle factors

Lifestyle factors can have a profound influence on human healthspan, lifespan and SCI status (Furman et al., 2019). To understand this better in the context of InflammAge, we tested the association of InflammAge acceleration with lifestyle factors such as self-reported smoking status, alcohol consumption, physical activity and different dietary variables in the Generation Scotland cohort.

Smoking status displayed a significant association with InflammAge acceleration (ANOVA p-value=1.1x10^-18^). The effect behaved in a dose-response manner, with non-smokers showing the lowest InflammAge acceleration, followed by ex-smokers (former smokers) and current smokers (ex-smoker vs non-smoker: Diff=-0.70, 95% CI [-0.95--0.44)], Tukey adj. p-value=3.2x10^-8^; smoker vs non-smoker: Diff=1.03, 95% CI [0.73-1.33], Tukey adj. p-value=3.2x10^-8^) (**Fig. 4a**). With regards to alcohol consumption, InflammAge acceleration was higher in heavy drinkers (≥14 units / week) compared to moderate drinkers (<14 units / week; Diff=0.62, 95% CI [0.30-0.95], Tukey adj. p-value=1.8x10^-5^), while lower in moderate drinkers compared to non-drinkers (Diff=-0.48, 95% CI [-0.83--0.12], Tukey adj. p-value=4.4x10^-3^) (**Fig. 4b**).

**Fig. 4.**
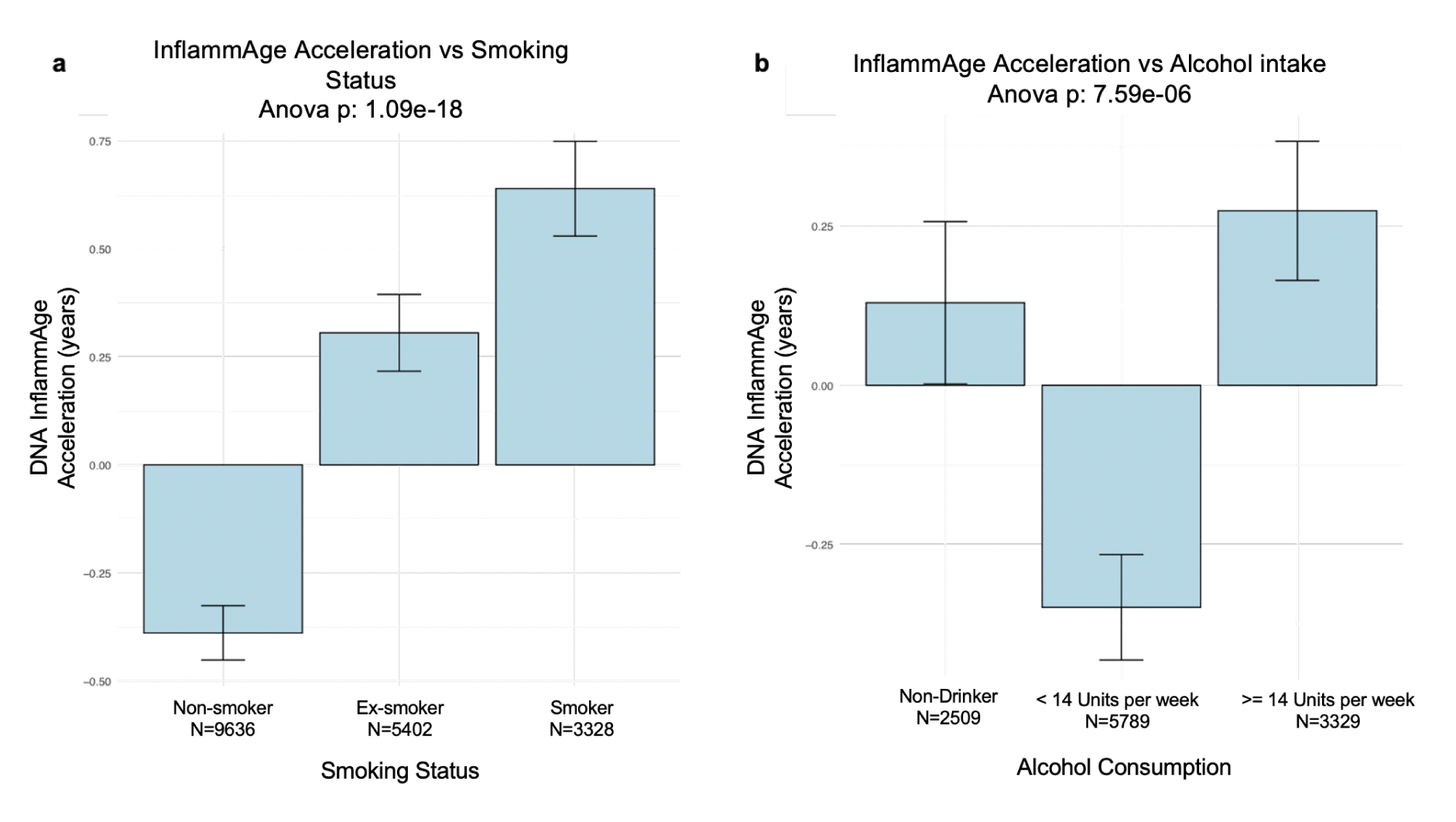
InflammAge acceleration associations with smoking status and alcohol intake in the Generation Scotland cohort. a. Barplot with ANOVA p-value for self-reported smoking status. **b** Barplot with ANOVA p-value for self-reported alcohol consumption in the week before the blood sample for DNAm measurement was taken. Plots show mean InflammAge acceleration per group and error bars show standard errors of the mean.

Self-reported physical activity (as measured in total metabolic equivalents (MET) minutes) was not associated with InflammAge acceleration (SCC=-0.01, p-value=0.16). This was unexpected, given the links between exercise and reductions in systemic chronic inflammation (Hayashino et al., 2014). However, this could be due to the nature of self-reported exercise data, which tends to be less accurate than wearable-based data (Beagle et al., 2020).

When analysing the association of dietary consumption patterns with InflammAge acceleration, we ran linear models adjusted for the same confounding factors used in the disease outcome analysis (**Table 1**, see Methods). An intake of oily fish at any frequency was significantly associated with lower InflammAge acceleration compared to never having oily fish (Effect size=-0.31, SE=0.13, p-value=0.02). Intake of other dietary constituents at any frequency compared to never showed no significant difference when adjusted for confounding factors. However, individual dose groups showed significance in the unadjusted ANOVA model (see **Supplementary Table S5)**. This included a lower InflammAge acceleration for those individuals with a higher intake of brown bread (5-6 days/week vs never: Diff=-0.94, 95% CI [-1.66--0.23], Tukey adj. p-value=2x10^-3^), higher fruit (daily vs. once per week: Diff=-0.84, 95% CI [-1.51--0.17], Tukey adj. p-value=4.4x10^-3^) and higher green vegetables intake (5-6 days/week vs never: Diff=-1.08, 95% CI [-2.00--0.17], Tukey adj. p-value=8.8x10^-3^). On the contrary, a higher intake of meat showed a trend towards higher InflammAge acceleration (intake daily vs < once per month: Diff=1.15, 95% CI [0.17-2.12], Tukey adj. p-value=9.7x10^-3^) although this was not observed across all dietary consumption amounts. Interestingly, these effects are consistent with the literature reported for other blood-based epigenetic clocks (Quach et al., 2017, Levine et al., 2018, Lu et al., 2019).

**Table 1.** Association analysis for InflammAge acceleration with dietary variables in Generation Scotland. Summary statistics for the covariate-adjusted linear models testing an association between InflammAge acceleration and different dietary variables (ever vs never categories, collected at baseline during GS participant recruitment). For the full summary statistics of all ANOVA comparisons see **Supplementary Table S5.**

| Dietary category | Effect size | Standard error | P-value |
| --- | --- | --- | --- |
| Oily fish | -0.305 | 0.134 | 0.023 |
| Dairy | -1.017 | 0.664 | 0.126 |
| Brown Bread | -0.315 | 0.216 | 0.145 |
| Other fish | 0.232 | 0.213 | 0.275 |
| Eggs | -0.223 | 0.358 | 0.534 |
| Liver | -0.053 | 0.105 | 0.613 |
| Green vegetables | -0.147 | 0.302 | 0.627 |
| Poultry | 0.120 | 0.269 | 0.656 |
| Other meat | 0.089 | 0.251 | 0.724 |
| Fruit | -0.150 | 0.470 | 0.749 |
| Other vegetables | -0.128 | 0.570 | 0.823 |

Overall, our results suggest that InflammAge has potential to monitor the impact of lifestyle factors on SCI.

### InflammAge recapitulates immunosenescent cell type composition in blood

To understand if InflammAge is linked to the relative abundance of different cell type populations, we performed cell type deconvolution in the blood DNAm data from Generation Scotland. We then calculated the correlation of InflammAge acceleration with the estimated proportions of each cell type in the dataset. InflammAge acceleration showed significant associations with predicted immune cell type proportions in blood despite not using EpiScores related to cell type deconvolution as part of training the model. InflammAge acceleration was positively correlated with monocytes (PCC=0.15), neutrophils (PCC=0.44) and CD8+CD28-CD45RA- T cells (memory/effector CD8+ T cells, PCC=0.26) (Tomiyama et al., 2002). In contrast, InflammAge acceleration was negatively correlated with B cells (PCC=-0.20), CD4+ T cells (PCC=-0.47), total CD8+ T cells (PCC=-0.121) and naïve CD8+ T cells (PCC=-0.24). Eosinophils (PCC=-0.02) and NK cells (PCC=-0.05) showed almost no change with InflammAge acceleration (**Fig. 5**). This is consistent with changes observed in the immune system during human ageing and immunosenescence (Koestler et al., 2017; Martín-Herranz, 2019; Saule et al., 2006). Thus, individuals that have positive InflammAge acceleration (InflammAge results higher than expected) display a more immunosenescent profile.

**Fig. 5.**
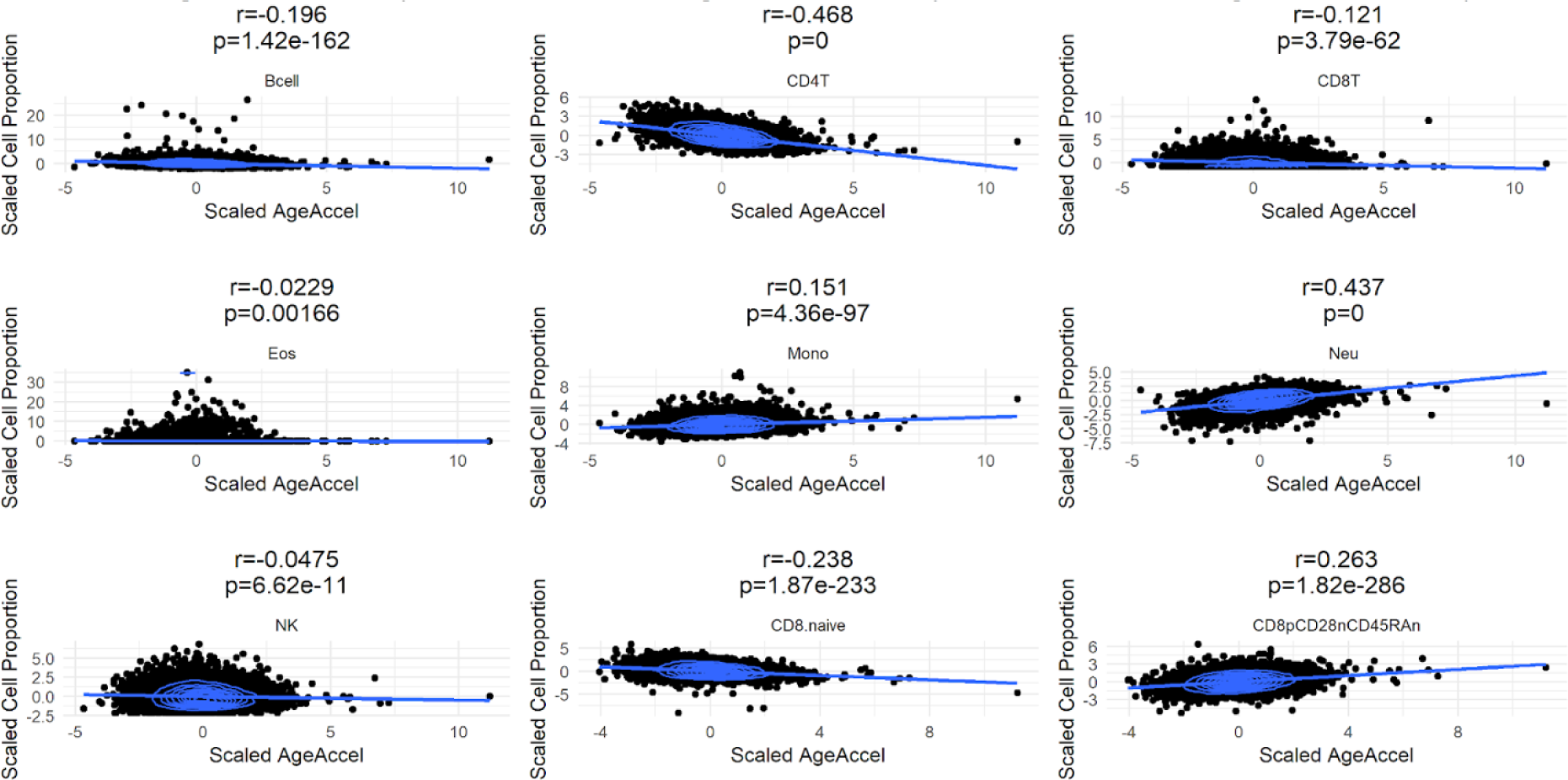
InflammAge acceleration recapitulates immunosenescent cell type composition. Scatterplot of InflammAge acceleration (x-axis) and blood cell type proportions (y-axis) estimated from blood DNAm data in Generation Scotland. Values were scaled to be centred around zero (SD = 1). Different scatterplots represent the different cell types. Abbreviations: Bcell = B-cell, CD4T = CD4+ T-cells, CD8T = CD8+ T-cells, Eos = eosinophils, Mono = monocytes, Neu = neutrophils, NK = natural killer cells, CD8.naive = naive CD8+ T-cells, CD8pCD28nCD45RAn = memory/effector CD8+ T-cells, r = Pearson correlation coefficient, p = p-value.

## 3. Discussion

Preventative healthcare and interventions that target ageing directly can have a profound health and economic impact on our societies. A slowdown in ageing that results in an increase in human life expectancy by 1 year is understood to be worth US$38 trillion (Scott et al., 2021). This can be transformational for many healthcare systems around the world and patients who currently live carrying the burden of multi-morbidity. In order to understand the ageing process and the risk of developing age-related diseases, accessible and actionable biomarkers need to be developed and made widely available.

In this work, we have created a framework that facilitates the creation of novel saliva-based DNA methylation biomarkers. Saliva is a very accessible and non-invasive sample type. Saliva cell composition comprises ∼80% immune cells and ∼20% epithelial buccal cells (Zheng et al., 2018). This cell composition similarity likely results in strong correlations (*r*∼0.97) between saliva and blood DNAm patterns (Nishitani et al., 2023). This is higher than the blood-saliva correlations typically observed when directly measuring inflammatory markers (such as salivary CRP) (Szabo & Slavish, 2021). All of this supports the case for saliva DNAm as a clinically meaningful and reproducible data type to build biomarkers, and this is indeed what we observe in this study (with InflammAge ICC > 0.8 within saliva). Given that most biological data in humans, including DNAm data, has been generated in blood samples, the ability to transfer biological information from blood to saliva DNAm opens the door for preventative healthcare and population screening applications at scale, including at-home testing and reaching isolated populations. This has also been explored by other authors using different methodologies (Galkin et al., 2021).

We have illustrated the value of our biomarker discovery framework by creating InflammAge, a novel saliva-based DNAm biomarker for systemic chronic inflammation (SCI), one of the hallmarks of ageing. InflammAge was built bottom-up by selecting DNAm features that are associated with chronic inflammatory markers, which allows for biological interpretability of the results, unlike several existing epigenetic clocks. This is critical given that different individuals may age differently, i.e. have different “ageotypes”, thus, personalised interventions need to recognise this (Ahadi et al., 2020; Oh et al., 2023). Improved biological interpretability may also be helpful in clinical trials to improve patient stratification and even increase understanding of the mechanism of action of compounds targeting ageing related pathways, thus serving as a ‘discovery biomarker’ (Moqri et al., 2023). Using DNAm features that aggregate different CpG probes, we also seem to benefit from dimensionality reduction as a de-noising mechanism, which has been reported to significantly improve the reliability of epigenetic clocks (Higgins-Chen et al., 2022).

During feature selection for InflammAge, we uncover DNAm predictors of blood inflammatory proteins that change consistently during ageing. Importantly, many of these EpiScores were trained on protein levels adjusted for age, sex, cohort-specific variables and potential genetic protein quantitative trait loci (pQTL) effects (Gadd et al., 2022); which requires careful interpretation when comparing them to direct protein concentration measurements and drawing biological conclusions from these features. Nevertheless, features that show higher levels with increased age contain several examples of pro-inflammatory proteins, such as IL-6, a classic biomarker for inflammageing (Ferrucci & Fabbri, 2018), TNFRSF1B, one of the receptors of TNF-alpha (Geurts et al., 2000), or CXCL10, a cytokine induced by TNF-α and IFN-γ (Vazirinejad et al., 2014). The top feature positively associated with age is CHIT1 (chitinase 1 or chitotriosidase), an enzyme produced and secreted by activated macrophages to fight chitin-containing pathogens, such as fungi and house-dust mites (Mack et al., 2015). CHIT1 is dysregulated in a variety of diseases characterised by chronic inflammation, such as pulmonary fibrosis, bronchial asthma, COPD and pulmonary infections (Chang et al., 2020).

Monocytes and macrophages are considered key effector cellular populations for inflammageing. Traditionally, M1 macrophages are thought to be pro-inflammatory while M2 macrophages are anti-inflammatory and involved in tissue repair (De Maeyer & Chambers, 2021); although a recent study using single-cell technologies has shown that macrophage polarisation is more complex than the M1/M2 dichotomy (Cui et al., 2023). Interestingly, M1/M2 macrophage populations change during human ageing (Costantini et al., 2018) and we observe signals that InflammAge may be identifying this phenomenon. For example, there is an increase in the DNAm proxy for proteins like CCL18, which causes M2 macrophage maturation (Schraufstatter et al., 2012), or MMP12, a matrix metalloprotease secreted by M2 macrophages (Aristorena et al., 2019). Additionally, NOTCH1 is the top feature negatively associated with age. Notch signalling has been proposed as a target in chronic inflammatory diseases (Christopoulos et al., 2021) and its inhibition increases M2 macrophage polarisation (Wang et al., 2010). Further studies should look at this phenomenon in more detail.

In this work, we also uncover that higher InflammAge acceleration resembles immunosenescence in blood. A link between an increase in SCI and the decreased ability of the immune system to fight infection with age has been widely reported in the literature. Indeed, we observe an association between predisposition to several infections and a higher InflammAge acceleration in blood (see **Fig. 3a**). This link is complex, with some authors suggesting that higher SCI may actually be protective in certain elderly populations where endemic infectious disease is present (Batista et al., 2020) and even improve response to vaccination (Picard et al., 2022). InflammAge may be helpful to untangle this complex relationship in human populations. Moreover, given that InflammAge acceleration is associated with immunosenescent cell composition changes, future research should look at the impact of this effect for the biomarker (e.g. comparing the current ‘extrinsic’ InflammAge with a a cell-composition corrected or ‘intrinsic’ version of InflammAge) (Martin-Herranz et al., 2019).

Thanks to the rich electronic health record data available in Generation Scotland, we have performed, to the best of our knowledge, the most comprehensive screening to assess the ability of a new epigenetic clock to predict clinical outcomes. We have demonstrated that InflammAge captures associations with all-cause mortality, different disease outcomes and lifestyle factors; in many cases outperforming CRP, a widely-used gold standard SCI biomarker. Interestingly, this seems to be the case for most of the cancer outcomes. For example, InflammAge shows a higher hazard ratio for non-Hodgkin’s lymphoma (NHL) when compared to CRP. A meta-analysis of 17 studies identified that elevated levels of several inflammatory biomarkers (such as TNF-α) are associated with a higher risk of developing NHL, potentially implicating them in the aetiology of the disease (Makgoeng et al., 2018). Albeit some of the model assumptions were violated, leukaemia is another cancer type where InflammAge improves results from CRP. A recent single-cell multi-omics analysis identified a role for chronic inflammation as a driver of leukemic evolution (Rodriguez-Meira et al., 2023). Other epigenetic clocks are generally powerful to capture associations with cancer (Lu et al., 2019), and we also observed this for the skin-blood clock in our data (which otherwise performs poorly across the rest of disease categories). InflammAge also tends to outperform CRP in the disease categories “haematological/immunological conditions” and “infectious diseases”. The latter could be reflective of the ability of InflammAge to quantify immunosenescence (previously discussed). Interestingly, while CRP is better at capturing all-cause mortality in the short term (2 years), InflammAge matches CRP in the longer term (16 years+), perhaps indicating that InflammAge captures the medium to long-term risk of disease (as opposed to acute risk).

This study has several limitations. First, the saliva DNAm training dataset was relatively small (N=338). As more saliva DNAm data from consenting individuals becomes available, the performance of our biomarker discovery framework and new versions of InflammAge are likely to improve. Furthermore, the current saliva DNAm datasets do not have linked mortality or disease outcome data, which precluded us from a saliva-only training and testing set-up, necessitating training on chronological age and testing in a large blood DNAm dataset like Generation Scotland instead. This also means that a direct comparison against other blood-trained epigenetic clocks (like PhenoAge, DunedinPACE or GrimAge) would be biased in favour of blood-based epigenetic clocks as the test dataset is blood. To perform that comparison and understand better the pros and cons of InflammAge, a blood-based InflammAge biomarker would allow for a more direct comparison. Additionally, sex differences were observed for our biomarker, with the male population displaying a higher InflammAge acceleration. Given that organs and systems display sex-specific ageing processes (including immune system ageing) (Márquez et al., 2020), investigating this in more detail (or even training sex-specific biomarkers) is necessary. When looking at the proposed extension of the NIH-FDA Biomarkers EndpointS and other Tools (BEST) classification framework recently suggested by the ‘Biomarkers of Ageing Consortium’ (Moqri et al., 2023), InflammAge is best suited to be classified as a predictive or prognostic biomarker. Understanding how InflammAge changes longitudinally and how responsive it is upon intervention will be important to assess in future studies.

Finally, we want to explore if our saliva-based DNAm framework can be used to deconvolute the ageing process and quantify many of the hallmarks of ageing into accessible biomarkers that are biologically interpretable. InflammAge shows a promise into this possibility, which can be generalised to quantify other complex phenotypes or risk factors beyond systemic chronic inflammation. We envision a saliva-based and cost-effective DNAm test that is able to personalise follow-up diagnostics and interventions and that becomes the entry point for a preventative healthcare journey for the global asymptomatic population.

## 4. Materials & Methods

### Hurdle DNAm platform

To create the Hurdle DNAm platform, we compiled a database of CpG probes reported to be associated with phenotypes relevant to health and disease as well as actionability. This was done using public resources that have collated genome-wide DNAm EWAS results, namely the EWAS catalogue (Battram et al., 2022) and EWAS Atlas (Li et al., 2019) (both accessed March 2022). Both resources were harmonised to remove outliers and errors. To obtain a list of genome-wide significant associations, we then filtered all CpG probe associations by p-value with a cutoff of p<3.6x10^-8^ as recommended for the Illumina Infinium

Methylation450 BeadChip array (Saffari et al., 2018). To form a core set of highest confidence EWAS associations, we intersected studies from both resources and included up to 1,000 CpG probes per study (**Supplementary Fig. S1a**). We further manually curated papers and preprints reporting EWAS results for phenotypes of interest and included CpGs associated with different epigenetic clocks, cell type composition, and epigenetic proxies for blood proteins, adding a further 47 publications. In total we included >30,000 CpG probes reported in 212 publications (**Supplementary Fig. S1b**). These CpG probes were deployed in a cost-effective Illumina methylation array that we named the Hurdle DNAm platform.

### Training dataset

We created a training dataset with saliva genome-wide DNA methylation (DNAm) samples by combining (N=338, **Supplementary Table S1**): a) published datasets; including datasets from GEO (Edgar et al., 2002) and ArrayExpress (Parkinson et al., 2007); all measured on the Illumina Infinium Methylation450 BeadChip array); b) in-house saliva DNAm samples; from individuals based in the UK who provided informed consent for their anonymised data being used for research purposes; all measured on the Illumina Infinium MethylationEPIC v1.0 BeadChip array.

Chronological age and sex were also available for individuals in the dataset and only data from participants >=18 years old were used. Where cohorts indicated the disease status of participants, only samples from healthy controls were used. The resulting training dataset was used for validating the correlation between SCI-related EpiScore features with chronological age in saliva and also for training the final InflammAge model.

### Reliability dataset

Data used for reliability analyses was generated as part of an independent study with healthy older adults aged ≥60 years (manuscript in preparation, Prof. Lord personal communication). The study was approved by the UK Health Research Authority and the London-Bromley Research Ethics Committee (Reference: 22/PR/0698). The study was conducted in accordance with the Declaration of Helsinki, with informed and written consent obtained from all participants prior to recruitment into the study. Saliva and venous blood samples of 83 adults were collected at two time points with age 72.24 ± 6.46 years; BMI: 26.62 ± 3.99 kg/m^2^; 31 males, 52 females (a full design of the DNAm data generated for this work can be found in **Supplementary Fig. S8**). At baseline (visit 1, V1) and week 12 (visit 2, V2), peripheral venous blood samples were acquired by venipuncture into vacutainers (BD Biosciences, Oxford, UK) containing ethylenediaminetetraacetic acid (EDTA) and stored at -80°C until DNA extraction. Saliva samples were collected at baseline and week 12 for downstream DNA methylation analysis using Hurdle’s sample collection kit (ORAgene DNA Saliva Collection kit; OG-600; DNA Genotek) and stored at 4°C prior to analysis. Genomic DNA from both saliva and whole blood were extracted and processed at Eurofins Genomics, Denmark. Bisulfite conversion was performed and samples were split into two and processed in parallel on the Illumina Infinium MethylationEPIC v1.0 BeadChip array (Illumina, CA, USA) and the Hurdle DNAm platform, creating technical replicates used for ICC calculation. All DNAm data were analysed using the same DNA methylation data processing pipeline (see below). We did not correct for array position or other technical factors as samples were randomised across arrays within each tissue and run with the same reagent batches.

### Test dataset: Alpha cohort

A dataset with matched blood SCI biomarkers and saliva DNAm was collected as part of a Hurdle human study. A cohort of 78 participants between ages 18-80 were recruited from a pool of 192 following exclusion criteria, which was further reduced to 61 participants after quality control for DNAm and blood results. Exclusion criteria were: acute infection in the past 6 weeks (including COVID-19 or influenza), prescription of immunosuppressant drugs, diagnosed chronic diseases affecting inflammatory status, heavy smokers (>20/day), harmful drinking habits (>35 units/week for women, >50 units/week for men as defined in (Gordon, 2001)), heavy exercise or significant diet changes prior to sampling. All individuals were based in the UK and provided informed consent for their pseudoanonymised data being used for research purposes. Saliva and venous blood samples were collected in the morning of the same day. Saliva samples were self-collected at home using Hurdle’s sample collection kit (ORAgene DNA Saliva Collection kit; OG-600; DNA Genotek). Peripheral venous blood samples were acquired by venipuncture in-clinic into vacutainers (BD Biosciences, Oxford, UK) containing ethylenediaminetetraacetic acid (EDTA), lithium heparin (LH) or z-serum clotting activator. Serum was isolated from vacutainers containing z-serum clotting factor by incubation at room temperature for 30 minutes to allow sufficient clotting, followed by centrifugation at 1620 x g for 10 minutes at room temperature, before storage at -80°C. Plasma was isolated from blood collected in vacutainers containing LH or EDTA by centrifugation at 461 x g for 8 minutes at room temperature. Plasma was stored at -80°C. DNA methylation was measured from saliva samples using the Illumina Infinium MethylationEPIC v1.0 BeadChip array (Illumina, CA, USA) at Eurofins Genomics, Denmark. Blood cytokine levels were quantified at Affinity Labs, London, UK. The following analytes were quantified in serum: albumin, complement C4, hsCRP (Siemens Advia 1800), and cortisol (Siemens Advia Centaur XP). The following analytes were quantified in plasma: IL-6, IL-1β, TNF-α, IFN-γ, IL-10, IL-8/CCL-8, IL-17, SAA (Mesoscale Discovery V Plex assay), CXCL-9, CXCL-1/GROα, GDF-15, TNFSF10/TRAIL, PAPP-A, CCL-11 (Biotechne ELISA), HbA1c, Full Blood Count: WBC, NEUT, MONO, LYM, ESO, BASO, RBC, HB, HCT, MCV, MCH, MCHC, RDW, PLT, MPV (Siemens Advia 560).

### Test dataset: Generation Scotland (GS) cohort

Generation Scotland is a family-based cohort consisting of individuals aged 18 to 99 years, living across Scotland. Participants were initially recruited from individuals registered at GP surgeries and then asked to invite first degree relatives to join the study, resulting in a final sample size of 23,960 individuals. For these participants, genetic as well as clinical, lifestyle, and sociodemographic information are available. A full description of the cohort and its recruitment has been reported previously (Smith et al., 2013).

DNA methylation was profiled in four sets between 2016 and 2022 using the Illumina Infinium MethylationEPIC v1.0 BeadChip array (Illumina, CA, USA). For all sets, the IDAT files were read into R using functions within *minfi* v.1.20.2 – 1.42.0 (Aryee et al., 2014). Quality control (QC) was applied to each set separately, at the time it was produced. Briefly, outliers were removed based on visual inspection of a plot of the log median intensity of the methylated versus unmethylated signal for each array (set 1). *ShinyMethyl*’s control probe plots were then inspected to identify outliers. Next, samples for which the sex predicted from the methylation data (based on the difference between the median copy number intensity for the Y chromosome and the median copy number intensity for the X chromosome) did not match the recorded sex were removed, with multi-dimensional scaling (MDS) plots generated and inspected for any additional sample outliers. The *pfilter* function within *wateRmelon* v.1.18.0 (Pidsley et al., 2013) was used to remove: (i) samples where ≥ 1% of CpGs had a detection *p*-value > 0.05; (ii) probes with a beadcount of < 3 in > 5% samples; and (iii) probes for which ≥ 0.5% of samples had a detection *p*-value > 0.05. QC of the DNA methylation data produced in sets 2-4 was carried out using both *Meffil* vs.1.1.0 and 1.1.2 (Min et al., 2018) and *shinyMethyl* versions 1.14.0 and 1.30.0 (Fortin et al., 2014). *Meffil* was used to perform dye-bias and background correction using the “noob” method (Triche et al., 2013). Samples were excluded if they were 1) affected by a strong dye bias or issues affecting bisulphite conversion (using default thresholds); 2) had median methylated signal intensity greater than three standard deviations lower than expected; or 3) had a different methylation-predicted sex to the self-reported sex. Deviations between methylation-predicted sex and self-reported sex were also assessed using *shinyMethyl*’s sex prediction function, which uses a different methodology to *Meffil*. *ShinyMethyl* was additionally used to plot the output of all control probes to permit the detection of outliers by visual inspection. Following these sample removal steps, *Meffil* was used to filter poor-performing samples and probes. Samples were removed if they: had > 0.5% CpG sites with a detection *p*-value > 0.01. Once the poor-performing samples had been removed, *Meffil* was re-run on the remaining dataset to identify poor-performing probes. These were defined as probes with a beadcount of < 3 in > 5% samples and/or > 1% samples with a detection *p*-value > 0.01. Quality-controlled DNA methylation data from the four datasets were combined and jointly normalised using the *dasen* method (Pidsley et al., 2013) (N_Set1_=5,055, N_Set2_=458, N_Set3_=4,374, N_Set4_=8,978).

DNAm InflammAge was calculated and mapped to phenotypic and clinical outcome information for 18,865 volunteers in Generation Scotland (GS). Clinical outcomes were obtained from linkage to primary care (GP) data and secondary care (General Acute Inpatient and Day Case) electronic health recrods from the Scottish Morbidity Record 01 (SMR01).

### DNA methylation data processing pipeline

This applies to all the DNA methylation data presented in this manuscript with the exception of the Generation Scotland cohort. Data processing was performed using RStudio (Version: 2023.09.1+494). IDAT files were processed using a custom pipeline that leveraged the *minfi* (Aryee et al., 2014) and *SeSaMe* (Zhou et al., 2018) packages. The pipeline uses *noob* background correction (Triche et al., 2013) and BMIQ within-array normalisation (Teschendorff et al., 2013) as previously described (Martin-Herranz et al., 2019). We removed cross-reactive probes, probes overlapping with SNPs and probes on sex chromosomes. Samples with overall call rates <70% or low mean intensity values were also removed from analyses. Depending on the analyses, we subset probes to a set of ∼30K included in the Hurdle DNAm platform before or after normalisation.

### Reliability analyses

We calculated intraclass correlation coefficients (ICC) using a customised version of the *dupicc* function of the *ENmix* R package version 1.36.08 (Xu & Taylor, 2021). We obtained results for a single-rater, absolute-agreement, two-way random-effects model (type=agreement, model = c(twoway), unit = single), in accordance with guidelines from Koo and Li (Koo & Li, 2016). For ICC calculation at the probe level, only probes present in both platforms (EPIC & Hurdle) after quality control were used (29,472). V1 samples of the reliability dataset were used for ICC calculations at the feature selection stage (steps 3 + 4 in the framework), and V2 samples were used for testing the reliability of the final InflammAge biomarker in step 7, therefore avoiding any data leakage. Comparisons between tissues and within tissues were performed with the same code. For EpiScore ICC assessment, EpiScores were calculated in each dataset and then used as input for the *dupicc* function. The same approach was taken for the InflammAge ICC calculation.

### Calculating EpiScores in saliva DNAm data

EpiScores were selected by literature review and calculated using the coefficients and methods provided in the original publications (Aslibekyan et al., 2018; Gadd et al., 2022; Hillary et al., 2020; Ligthart et al., 2016; Stevenson et al., 2020, 2021). In the case of the 109 EpiScores containing inflammatory proteins measured in the Olink and SOMAscan platforms (Gadd et al., 2022), the function *scale* (center=TRUE, scale=TRUE) from the *base* R package (R Core Team, 2023) was used to normalise the beta-values across samples in the dataset before calculating the EpiScores.

### EpiScore gene enrichment analysis

Gene Ontology (GO)-term enrichment was performed using the g:Profiler platform (Kolberg et al., 2023). Gene names of EpiScore proteins selected after ICC filtering were analysed in two sets, one with gene names of EpiScores positively associated with chronological age in saliva and one with negative association with age. All annotated genes were used as background for enrichment. Input gene names were ranked by using the absolute Pearson correlation coefficient with age in saliva. Enrichment results were visually inspected and top hits highlighted and saved.

### Training the InflammAge biomarker

Elastic net regression was the machine learning algorithm of choice to train a model that would predict chronological age (y_fit, dependent/output variable) using the selected SCI-related DNAm features (x_fit, independent variables). We used 10-fold cross-validation and alpha=0.5 as implemented in the *cv.glmnet* function in the *glmnet* R package (Friedman et al., 2010): cvfit_model <- cv.glmnet(x_fit, y_fit, alpha=0.5) We calculated age-adjusted InflammAge acceleration as the residuals from a linear regression model of DNAm InflammAge on chronological age (DNAm_InflammAge∼Chronological_age).

### Disease association analysis in Generation Scotland

A series of Cox proportional hazards regression models were run to relate InflammAge acceleration to the risk of developing 308 diseases, as coded by Kuan et al. (Kuan et al., 2023). Covariates included age at blood draw (when InflammAge was measured), sex, log-transformed BMI, years of education, socioeconomic status (Scottish Index of Multiple Deprivation), smoking pack years, and DNAm processing batch. Prior to the analyses, we calculated the proportion of females with an incident diagnosis for each disease. Where this was >90% (or <10%), we stratified the model to analyse females (males) only. Given that some diseases are more likely to occur at certain periods of life (e.g., dementia after the age of 65 years), we calculated the age of diagnosis for each disease and extracted the age of cases at the 2.5 and 97.5 percentiles. We then filtered the age at the event or censoring to fall within this window. This should retain a set of controls with similar profiles to the cases. Censoring was marked by being unaffected by the disease and either surviving to the latest point of linkage (April 2022) or time to death during follow-up. Regression assumptions were examined by tests of the Schoenfeld residuals, with p-value<0.05 being used to indicate a model violation. These analyses were repeated with additional biomarkers of SCI and ageing, for comparison with InflammAge acceleration, which included log-transformed measured CRP and skin-blood clock age acceleration (DNAmAgeSkinBloodClockAdjAge) (Horvath et al., 2018). These analyses only included diseases where at least 30 cases were present (to preserve participant anonymity and reduce the risk of models not converging).

In addition to the diseases listed in Kuan et al. (Kuan et al., 2023), we examined the relationship between InflammAge acceleration and a COVID-19 disease diagnosis, ascertained from primary and secondary care records. Cases were defined based on primary care Read codes 1JX1., 65PW1, A7951, 8CAO., 8CAO1, 9N312, and 4J3R1 (derived from Scottish Clinical Information Management in Practice coding (*System COVID-19 Codes – Available March 2020 Coding – Primary Care Informatics*, 2020), and secondary ICD10 codes U071 and U072. Logistic regression, adjusting for the same covariates as above, was used to analyse this relationship. Individuals who died without a prior COVID-19 diagnosis were removed prior to the analysis.

### Lifestyle factor association analysis in Generation Scotland

Alcohol consumption was binned into non-drinker (0 units in the last week), moderate drinker (<14 units in the last week) and heavy drinker groups (>=14 units in the last week). The sample was filtered to individuals who reported their last week’s alcohol intake as their usual weekly intake. Analysis of variance (ANOVA) tests were performed on smoking (non-smoker, ex-smoker, current smoker) and alcohol groups to first assess whether a significant relationship was present. Post-hoc Tukey tests were then performed to assess two-group relationships. Physical activity was reported as metabolic equivalents (MET) minutes and Spearman correlation of scaled MET minutes to scaled InflammAge acceleration was calculated.

Dietary intake data included self-reported information on frequency of brown bread, fruit, green vegetables, other vegetables, liver, poultry, other meat (non-poultry), eggs, dairy, oily fish, and other fish (non-oily). Two types of analyses were performed for dietary intake data:

An ‘ever/never’ diet variable was derived from each multi-level diet variable. Effect sizes and p-values were obtained from linear models that were adjusted with covariates previously described (**Table 1**): scale(InflammAgeAccel) ∼ age + sex + log(BMI) + education_years + scale(deprivation_rank) + smoking_status + DNAm_processing_batch + diet_variable In order to test associations between specific diet levels (doses), ANOVA and post-hoc Tukey tests were performed as previously described (**Supplementary Table S5**).

### Cell type composition analysis in Generation Scotland

For the Generation Scotland cohort (blood samples), the method and R function from Horvath 2013 was used (Horvath, 2013). Figures were plotted using *ggplot2* in R. Cell proportions were scaled to mean 0 and standard deviation 1 using the *scale* function.

Correlation between scaled DNAm InflammAge acceleration and scaled cell proportions was calculated for each cell type using Pearson correlation coefficient.

## 5. Data availability

Public training datasets (as listed in **Supplementary Table S1**) are available in public databases such as NCBI’s GEO (Edgar et al., 2002) and EMBL-EBI’s ArrayExpress (Parkinson et al., 2007). Data from Hurdle’s users or study participants (training, Alpha test, reliability dataset) cannot be made publicly available due to consent limitations, but requests for research collaboration can be submitted to the corresponding authors. Generation Scotland allows researchers to apply for access to their data through their collaboration proposal process. Further details can be found at https://www.ed.ac.uk/generation-scotland/for-researchers/access.

## Supporting information

Schmunk_et_al_bioRxiv_SupplementaryTables.xlsx

## Acknowledgements

We thank all study participants for their involvement. We further thank the following individuals for their support with this study: Ann Jordan, Evgenia Belova, Pavitra Chavan, Orla Codyre, Debora Crimella, Domenico Musto, Nabil El Nouby, Joanne Millar, Swetha Suresh, Harvey Watts, Ruth Atherton, Roughan Sheedy, Gordon McCulloch, Charles Ball, Felice Leung and Richard Rilstone. Generation Scotland received core support from the Chief Scientist Office of the Scottish Government Health Directorates [CZD/16/6] and the Scottish Funding Council [HR03006] and is currently supported by the Wellcome Trust [216767/Z/19/Z]. Genotyping of the GS:SFHS samples was carried out by the Genetics Core Laboratory at the Edinburgh Clinical Research Facility, University of Edinburgh, Scotland and was funded by the Medical Research Council UK and the Wellcome Trust (Wellcome Trust Strategic Award “STratifying Resilience and Depression Longitudinally” (STRADL) Reference 104036/Z/14/Z). The GS DNA methylation profiling and analysis was supported by Wellcome Investigator Award 220857/Z/20/Z and Grant 104036/Z/14/Z (PI: AM McIntosh) and through funding from NARSAD (Ref: 27404; awardee: Dr DM Howard) and the Royal College of Physicians of Edinburgh (Sim Fellowship; Awardee: Dr HC Whalley).

## 6. Author contributions

LJS, TPC, DLM, HJ, EZ and DEMH carried out the analyses. LJS, TPC, TMS and DEMH devised the study and secured the funding. KMG, JS, AC, AMM, TJ and JML performed study design and recruitment for the reliability dataset and contributed data. VJ, DK, NT, ES, MG, RT and the Hurdle bio-infrastructure team performed and supervised tech work and logistics. WJH, VO, KW, CEB and REM advised on the experimental and InflammAge validation strategy. WJH assisted in the generation of manuscript figures. LJS, TPC, HJ, TMS and DEMH wrote the first draft and all authors revised and approved the manuscript.

The Hurdle bio-infrastructure team includes: Pierric Descamps, Emily Taylor, Chuck Paiusi, Leona Mc Girr, Ollie Philpott, Ian Robinson, Hector Watson, Ana Magalia, Dave Russell and David Woods who are all employees at Chronomics Ltd., London, UK.

## 8. Supplementary Figures

**Supplementary Fig. S1.**
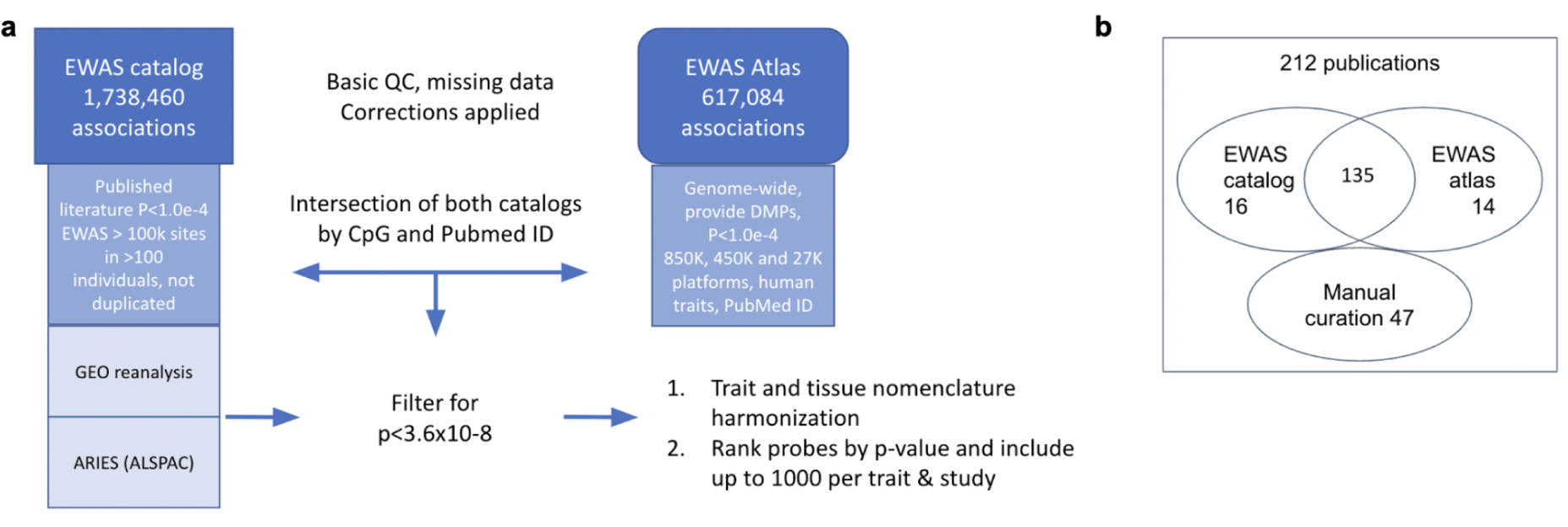
Hurdle DNAm platform design. a. Diagram showing how CpG sites were selected from the EWAS catalogue and the EWAS Atlas to be included in the Hurdle DNAm platform. **b** Summary of publication sources and numbers for the Hurdle DNAm platform design.

**Supplementary Fig. S2.**
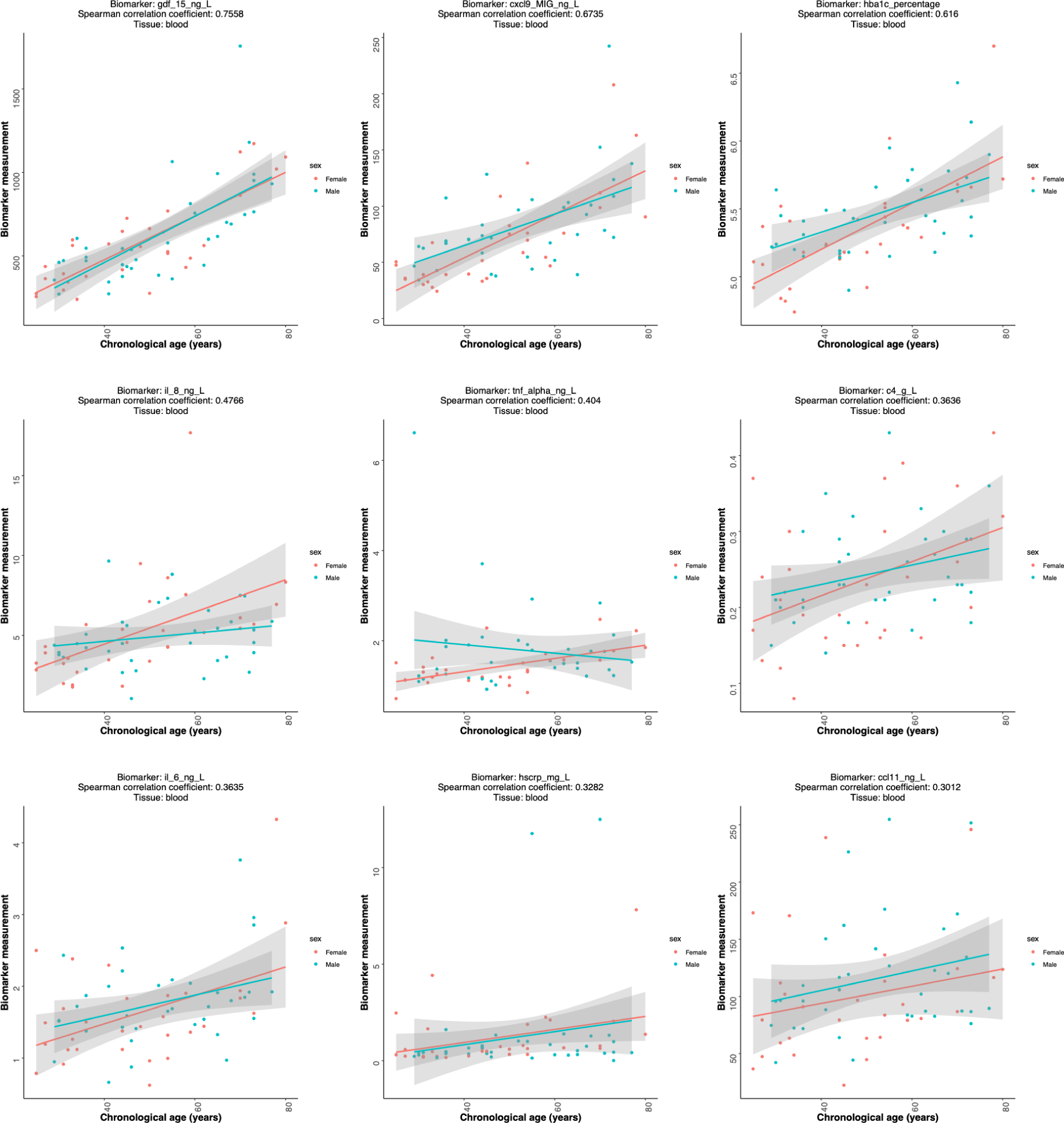
Blood biomarkers associations with chronological age in the Alpha test dataset. Scatterplots with Spearman correlation coefficients are shown for the association of 9 blood biomarkers measured from blood serum or plasma in the Alpha cohort study (see Methods). From top left to bottom right: GDF-15, CXCL9, HbA1c%, IL-8, TNF-α, C4, IL-6, CRP, CCL11.

**Supplementary Fig. S3.**
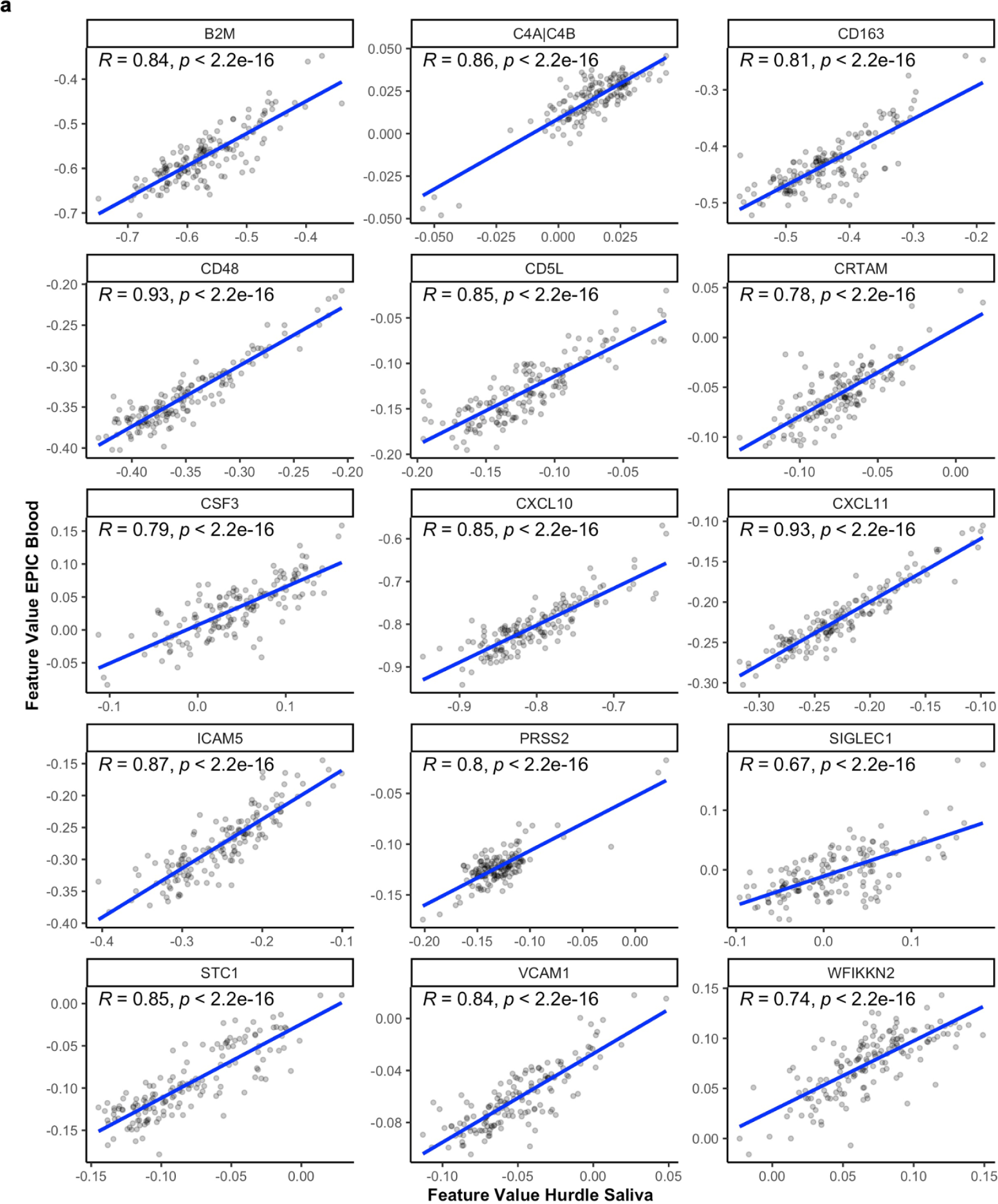
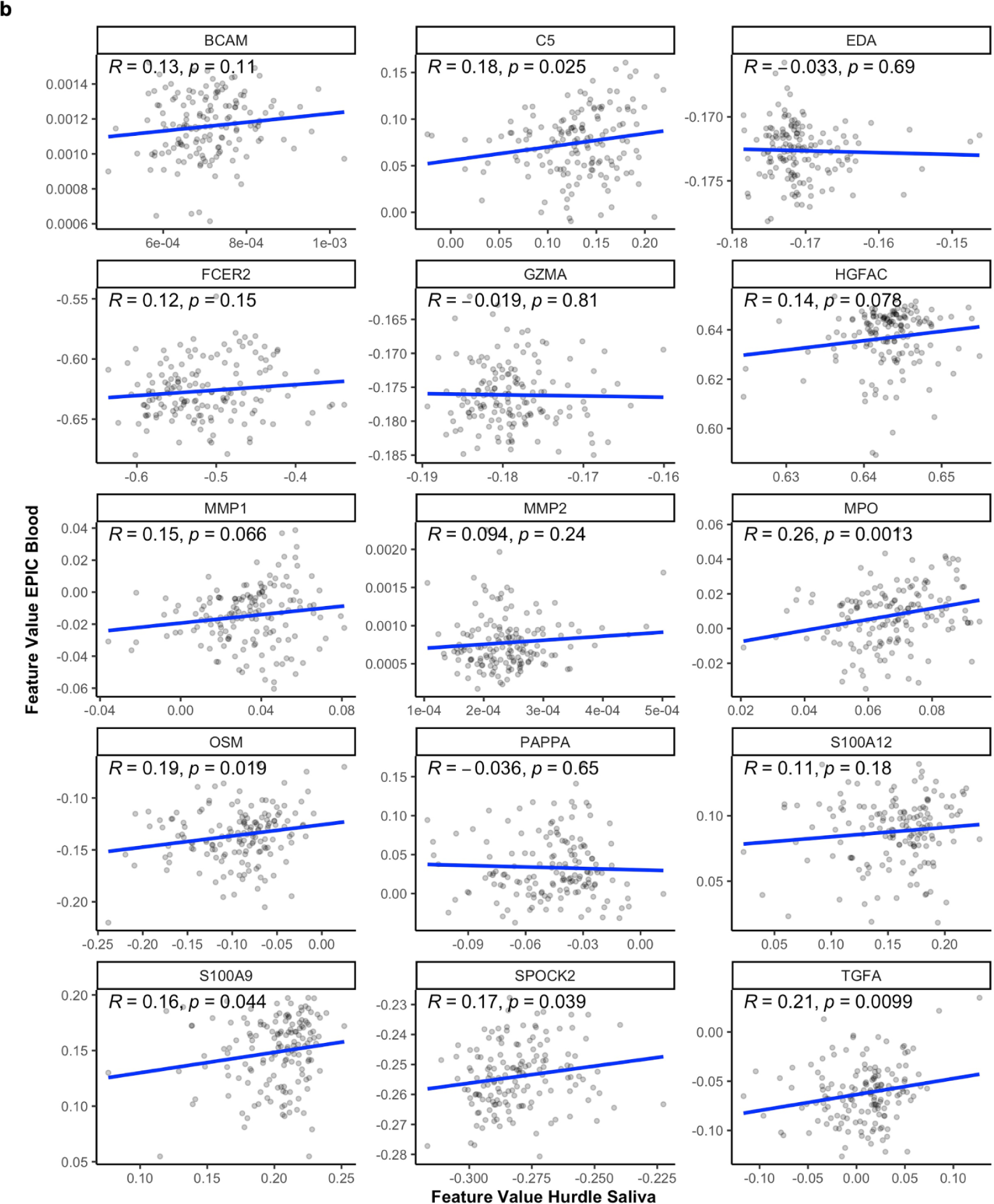
Cross-tissue correlations for EpiScore features in the reliability dataset. DNAm EpiScore values were calculated and compared between matched blood EPIC and saliva Hurdle samples. **a** Scatterplots with Pearson correlation coefficients (*R*) for the 15 EpiScores with the highest ICC values between blood EPIC and saliva Hurdle are shown. **b** Scatterplots with Pearson correlation coefficients (*R*) for the 15 EpiScores with the lowest ICC values between blood EPIC and saliva Hurdle are shown. *p* = unadjusted p-value for the Pearson correlation coefficient.

**Supplementary Fig. S4.**
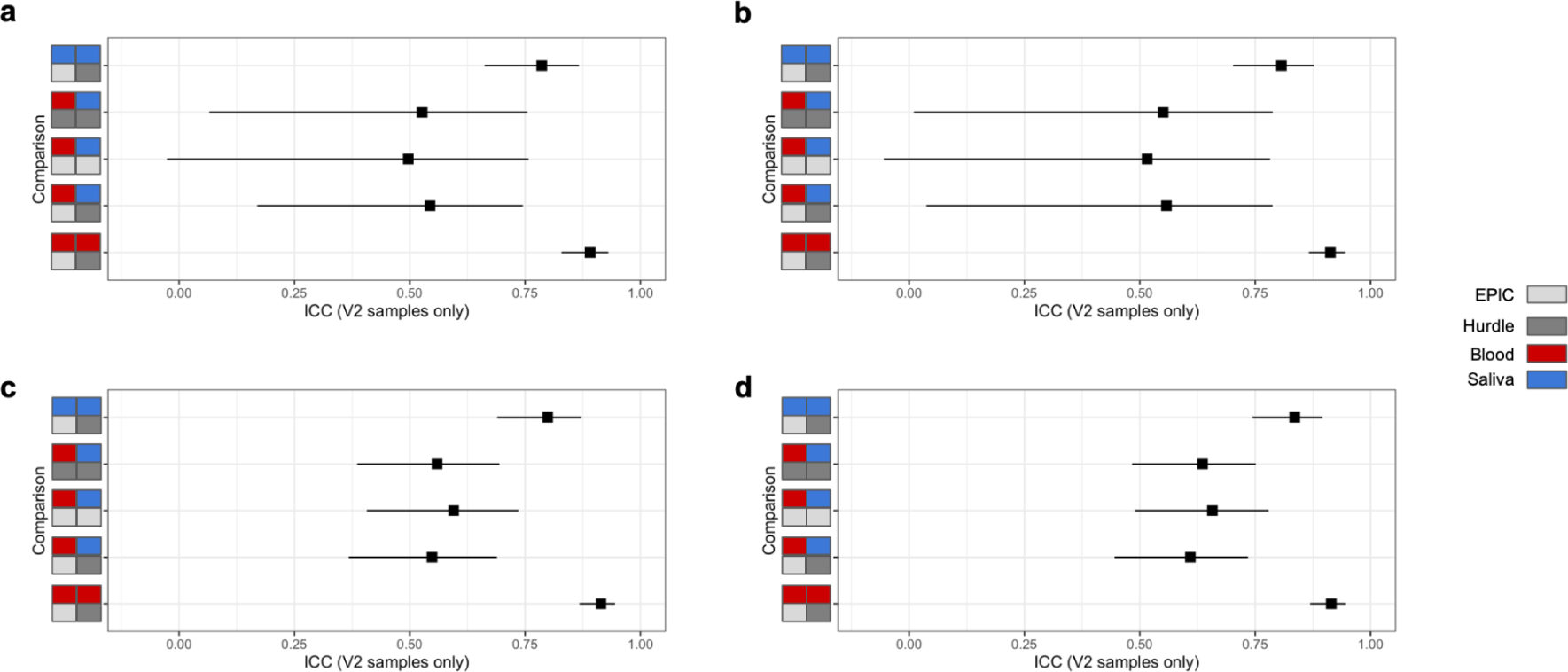
InflammAge ICC comparison across model versions. a. InflammAge trained with no EpiScore ICC filtering applied before training. **b** EpiScores filtered within tissues for icc_EbHb>0.75, icc_EsHs>0.75 (equivalent step 4). **c** EpiScores filtered within tissues for icc_EbHb>0.75, icc_EsHs>0.75, and across tissues for icc_EbEs>0.25, icc_HbHs>0.25 and icc_EbHs>0.25. **d** EpiScores filtered within tissues for icc_EbHb>0.75, icc_EsHs>0.75, icc_EbEs>0.10, icc_HbHs>0.10, icc_EbHs>0.10 (steps 3+4). E= EPIC, H= Hurdle DNAm platform, b = blood, s = saliva.

**Supplementary Fig. S5.**
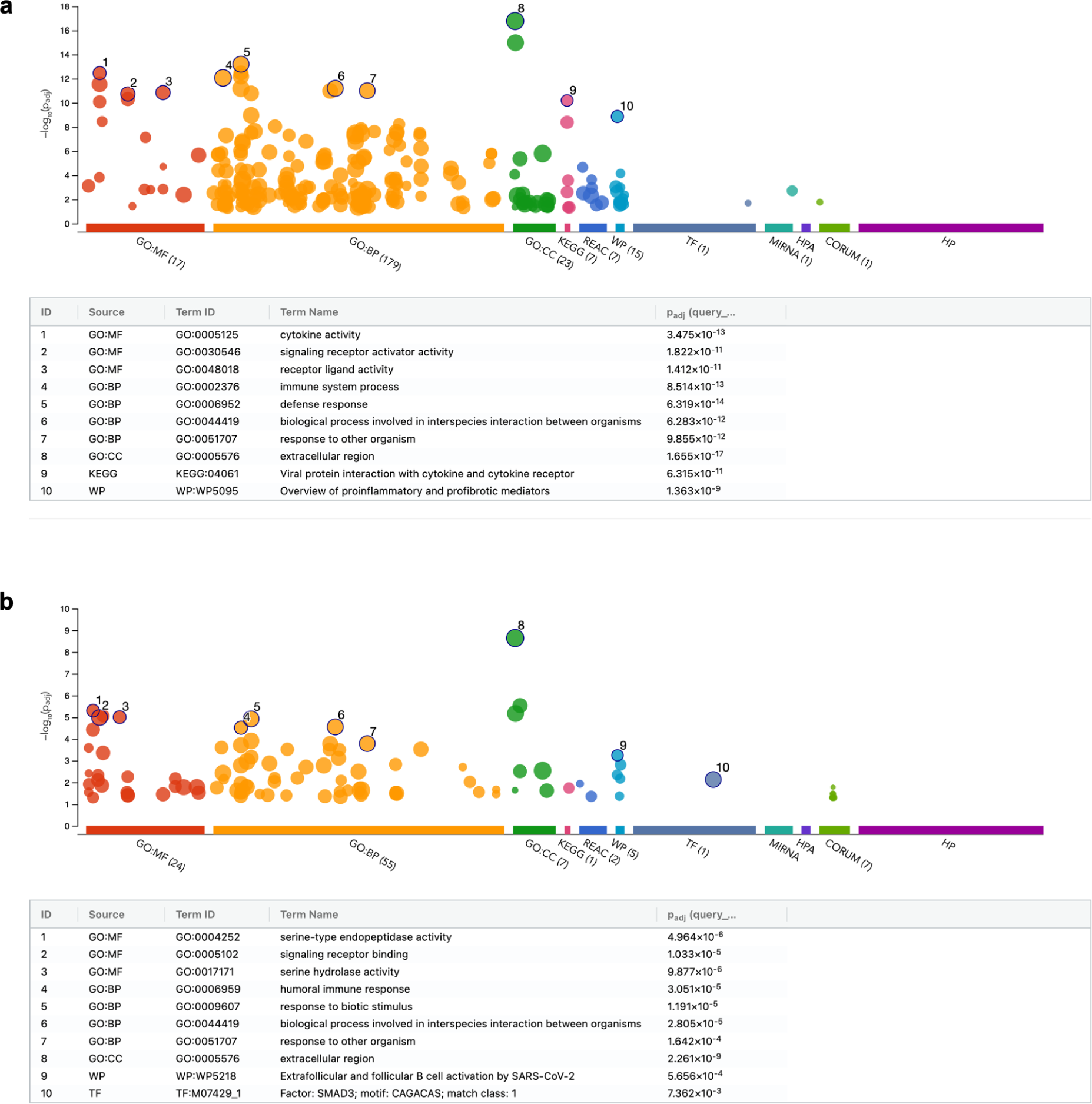
Gene ontology enrichment of EpiScore genes used as model training input. Generated as ordered queries using g:Profiler (Kolberg et al., 2023). **a** EpiScores with a positive association with chronological age in saliva were used as input (42). **b** EpiScores with a negative association with chronological age in saliva were used as input (21). Legends show 10 selected GO term names and adjusted p-values for enrichment against annotated genes.

**Supplementary Fig. S6.**
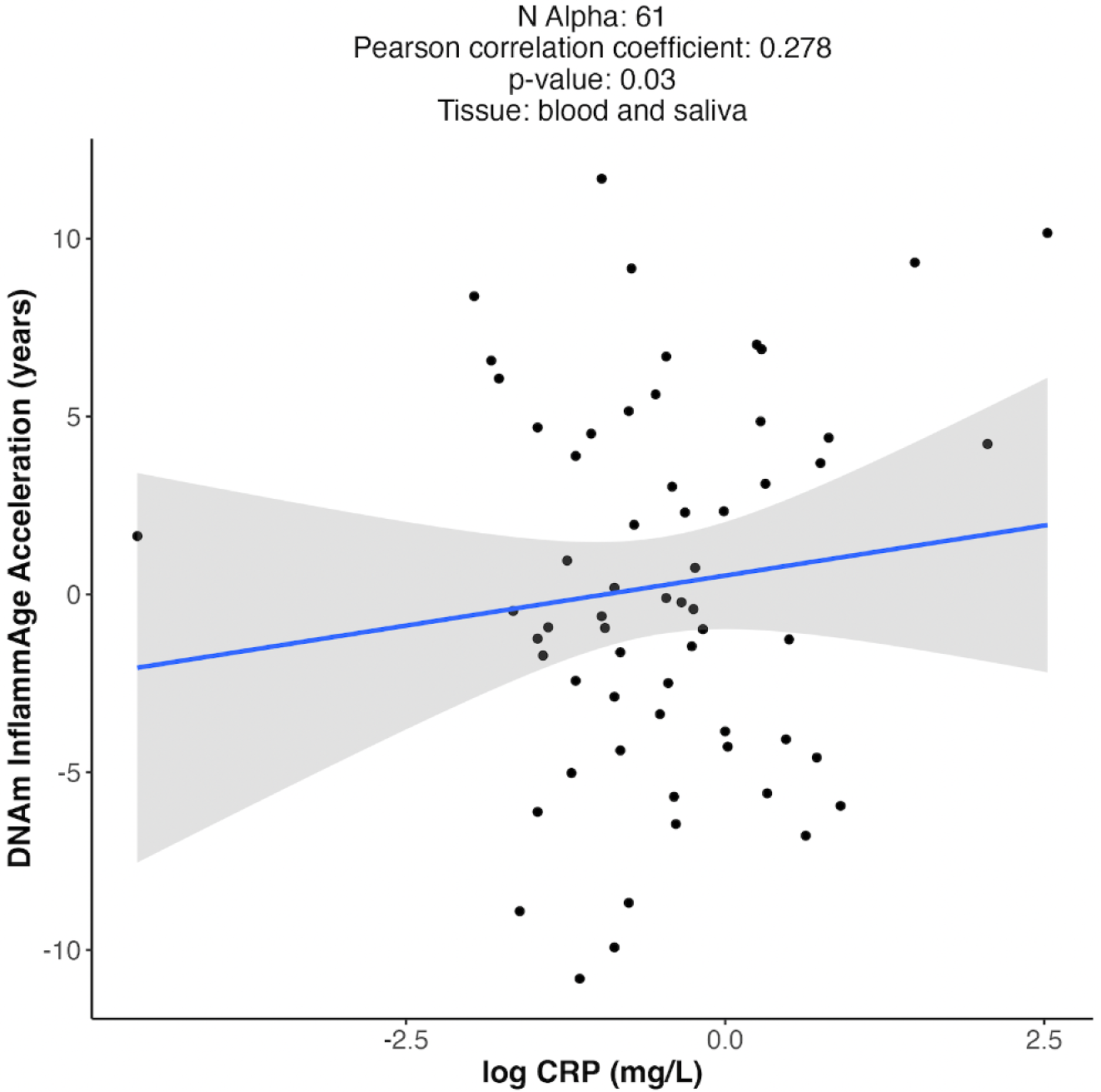
Scatterplot of InflammAge acceleration compared to log-transformed blood CRP in the Alpha test cohort. CRP was measured from blood serum (see Methods) and DNAm InflammAge acceleration was calculated from saliva DNAm data (N=61).

**Supplementary Fig. S7.**
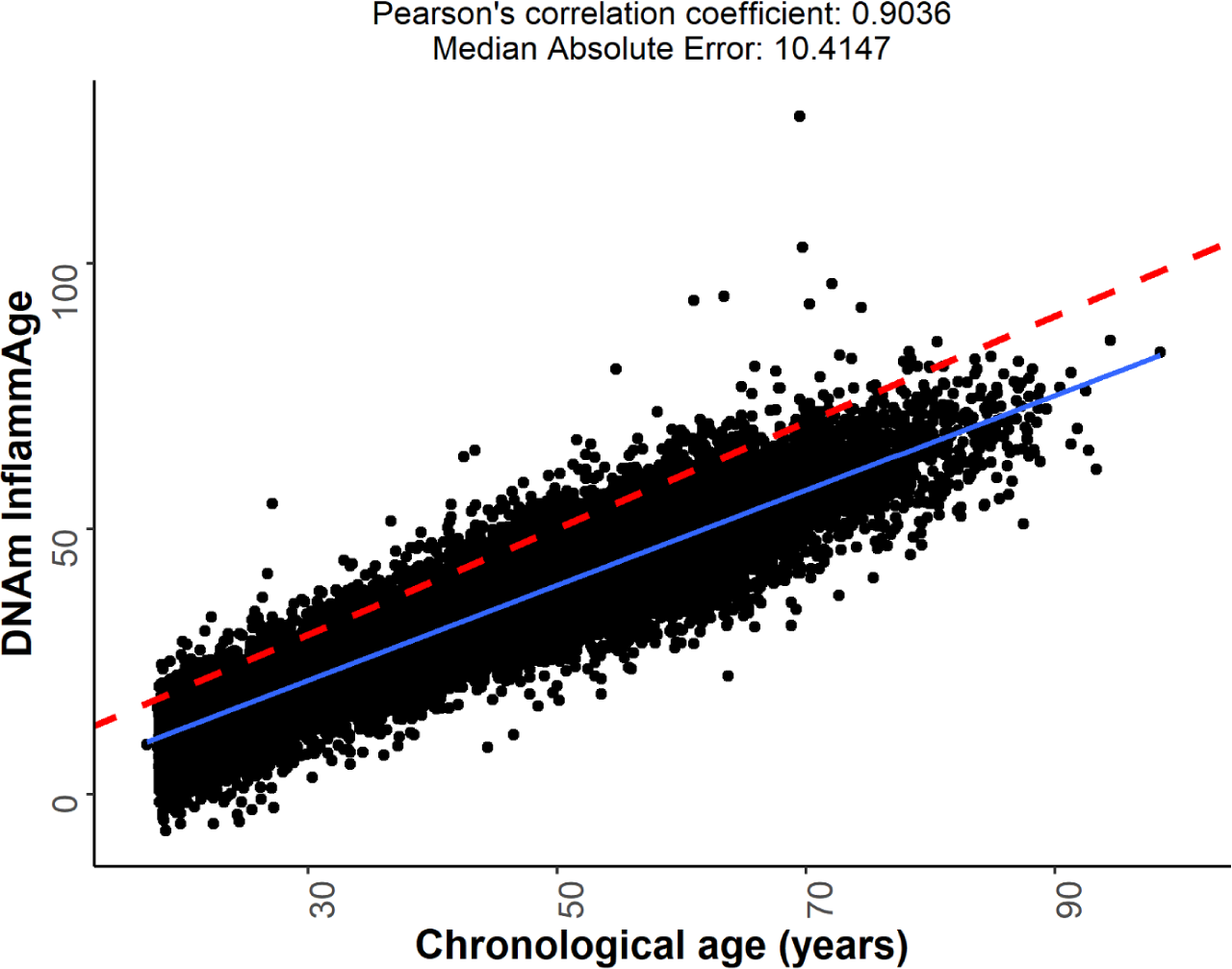
InflammAge performance in Generation Scotland. InflammAge was calculated for 18,865 participants blood DNAm data. The blue line shows the linear model fit; the red dashed line is the diagonal (intercept=0, slope=1).

**Supplement Fig. S8.**
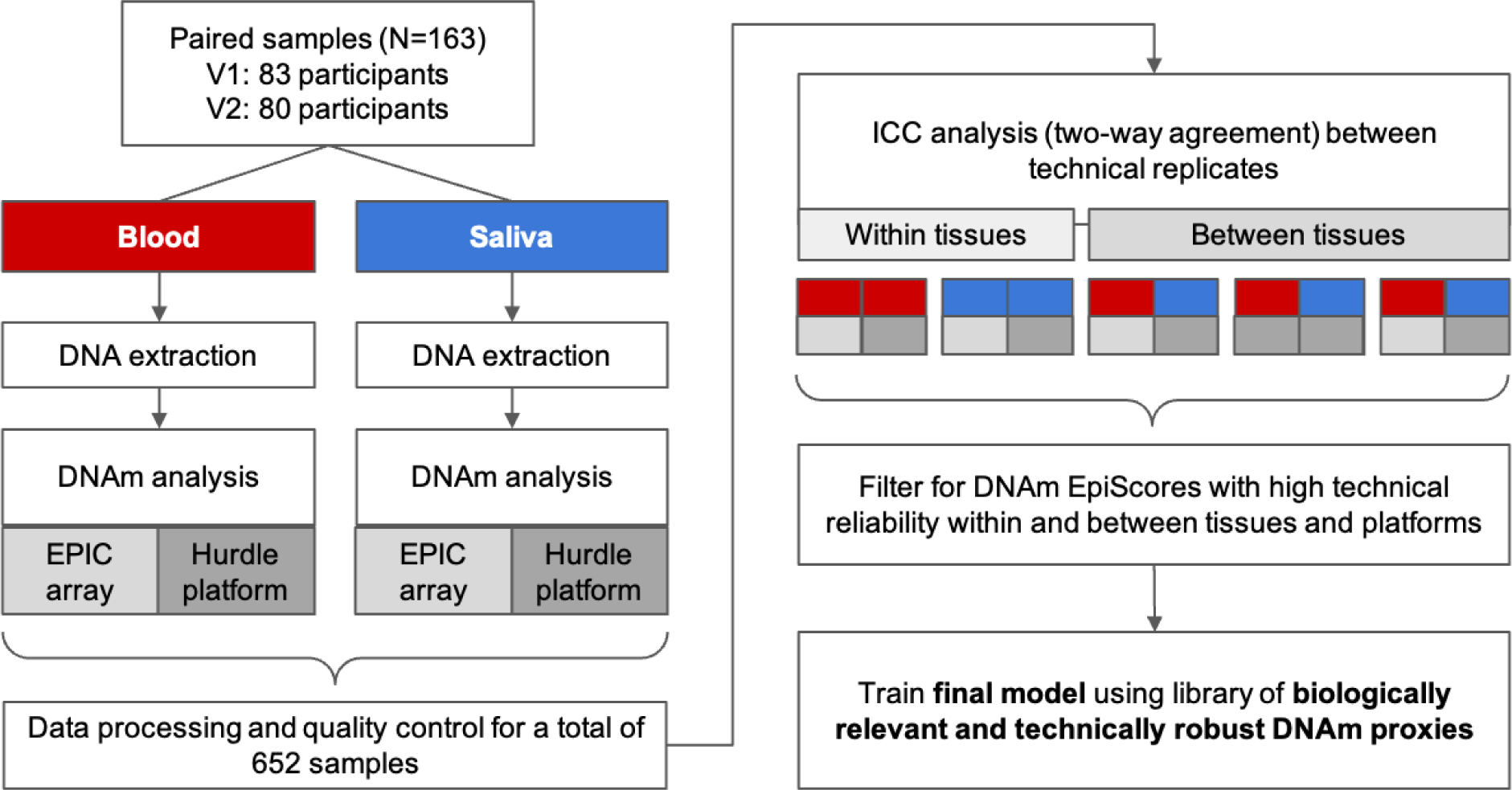
Reliability dataset ICC study design. A total of 83 participants were recruited and DNAm was measured in blood and saliva at baseline (V1, N=83) and after 3 months (V2, N=80). Samples were processed in duplicate on the EPIC (Illumina Infinium MethylationEPIC v1.0 BeadChip array) and the Hurdle DNAm platform and ICC was calculated for the five comparisons indicated on the right. ICCs calculated in V1 samples were used for filtering steps 3-4 of the framework. ICCs calculated in V2 samples were used to assess the reliability of the final InflammAge model in the training cohort.

## 9. Supplementary Tables

**Supplementary Table S1.** Overview of cohorts included in the InflammAge training dataset.

| Cohort | Platform | Tissue | Ethnicity | Disease status |
| --- | --- | --- | --- | --- |
| E-MTAB-5377 | Illumina Infinium Methylation450 BeadChip | Saliva | British | Control |
| GSE92767,<br>GSE59505,<br>GSE59508 |  | Saliva | Korean-Asian | Control |
| GSE111223-GSE78874 |  | Saliva | European + Hispanic | Control |
| GSE99029 |  | Saliva | South African Khomani San | Control |
| Hurdle in-house dataset | Illumina Infinium MethylationEPIC v1.0 BeadChip | Saliva | Not reported | Not reported |

**Supplementary Table S2.** Summary information for the Generation Scotland test cohort. It includes information for the risk factors that were included in the Cox regression models.

|  |  |
| --- | --- |
| **Total cohort size** | N = 18865 |
| **InflammAge Accel** |  |
| Mean (SD) | -0.000322 (6.32) |
| Median [Min, Max] | -0.106 [-29.5, 70.9] |
| **Age** |  |
| Mean (SD) | 47.6 (14.9) |
| Median [Min, Max] | 49.2 [17.1, 98.5] |
| **Sex** |  |
| Male | 7771 (41.2%) |
| Female | 11094 (58.8%) |
| **Smoking Status** |  |
| Non-smoker | 5402 (28.6%) |
| Ex-smoker | 9636 (51.1%) |
| Smoker | 3228 (17.1%) |
| Missing | 599 (3.2%) |
| **Socioeconomic status (SIMD)** |  |
| Mean (SD) | 3900 (1850) |
| Median [Min, Max] | 4340 [1.00, 6510] |
| Missing | 1153 (6.1%) |
| **Years of Education** |  |
| 0 | 7.00 (0.0%) |
| 1-4 | 50.0 (0.3%) |
| 5-9 | 546 (2.9%) |
| 10-11 | 4954 (26.3%) |
| 12-13 | 3888 (20.6%) |
| 14-15 | 2563 (13.6%) |
| 16-17 | 3525 (18.7%) |
| 18-19 | 1699 (9.0%) |
| 20-21 | 431 (2.3%) |
| 22-23 | 128 (0.7%) |
| 24 | 62.0 (0.3%) |
| Missing | 1012 (5.4%) |
| **Body Mass Index (kg/m2)** |  |
| Mean (SD) | 26.7 (5.18) |
| Median [Min, Max] | 25.9 [10.5, 71.4] |
| Missing | 120 (0.6%) |
| **Pack Years** |  |
| Mean (SD) | 8.30 (16.4) |
| Median [Min, Max] | 0 [0, 216] |
| Missing | 395 (2.1%) |

**Supplementary Table S3.**
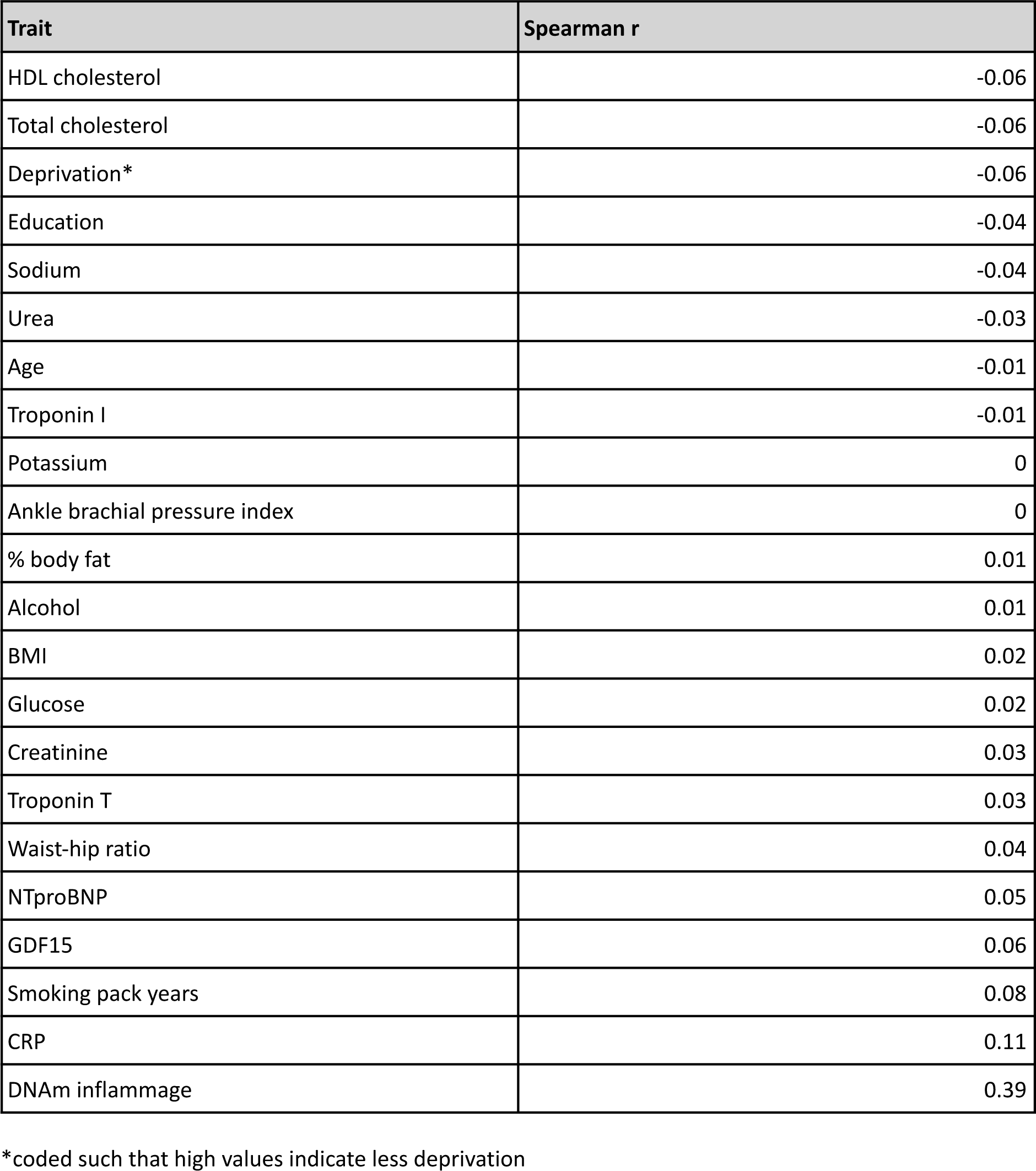
Spearman correlation coefficients between InflammAge acceleration and basic demographic and biochemistry variables in Generation Scotland (N=18,865).

**Supplementary Table S4. | Cox regression summary statistics across 65 diseases with >30 cases and 3 predictors (CRP, InflammAge acceleration, SkinBloodAge acceleration) in Generation Scotland.**

[Schmunk_et_al_bioRxiv_SupplementaryTables.xlsx]

**Supplementary Table S5. | Summary statistics across dietary lifestyle factors and InflammAge acceleration in Generation Scotland.** Tukey summary statistics for an association between InflammAge acceleration and different dietary variables collected at baseline during GS participant recruitment. **‘**diff’: mean difference in InflammAge acceleration between groups. ‘lwr’: lower 95% confidence interval. ‘upr’: upper 95% confidence interval. ‘p.adj’: Tukey p-value adjusted for multiple comparisons.

[Schmunk_et_al_bioRxiv_SupplementaryTables.xlsx]

## References

Ahadi, S., Zhou, W., Schüssler-Fiorenza Rose, S. M., Sailani, M. R., Contrepois, K., Avina, M., Ashland, M., Brunet, A., & Snyder, M. (2020). Personal aging markers and ageotypes revealed by deep longitudinal profiling. Nature Medicine, 26(1), 83–90. 10.1038/s41591-019-0719-5

Ambroa-Conde, A., Girón-Santamaría, L., Mosquera-Miguel, A., Phillips, C., Casares de Cal, M. A., Gómez-Tato, A., Álvarez-Dios, J., de la Puente, M., Ruiz-Ramírez, J., Lareu, M. V., & Freire-Aradas, A. (2022). Epigenetic age estimation in saliva and in buccal cells. Forensic Science International. Genetics, 61, 102770. 10.1016/j.fsigen.2022.102770

Aristorena, M., Gallardo-Vara, E., Vicen, M., de Las Casas-Engel, M., Ojeda-Fernandez, L., Nieto, C., Blanco, F. J., Valbuena-Diez, A. C., Botella, L. M., Nachtigal, P., Corbi, A. L., Colmenares, M., & Bernabeu, C. (2019). MMP-12, Secreted by Pro-Inflammatory Macrophages, Targets Endoglin in Human Macrophages and Endothelial Cells. International Journal of Molecular Sciences, 20(12). 10.3390/ijms20123107

Aryee, M. J., Jaffe, A. E., Corrada-Bravo, H., Ladd-Acosta, C., Feinberg, A. P., Hansen, K. D., & Irizarry, R. A. (2014). Minfi: A flexible and comprehensive Bioconductor package for the analysis of Infinium DNA methylation microarrays. Bioinformatics, 30(10), 1363–1369. 10.1093/bioinformatics/btu049

Aslibekyan, S., Agha, G., Colicino, E., Do, A. N., Lahti, J., Ligthart, S., Marioni, R. E., Marzi, C., Mendelson, M. M., Tanaka, T., Wielscher, M., Absher, D. M., Ferrucci, L., Franco, O. H., Gieger, C., Grallert, H., Hernandez, D., Huan, T., Iurato, S., … Arnett, D. K. (2018). Association of Methylation Signals With Incident Coronary Heart Disease in an Epigenome-Wide Assessment of Circulating Tumor Necrosis Factor α. JAMA Cardiology, 3(6), 463–472. 10.1001/jamacardio.2018.0510

Batista, M. A., Calvo-Fortes, F., Silveira-Nunes, G., Camatta, G. C., Speziali, E., Turroni, S., Teixeira-Carvalho, A., Martins-Filho, O. A., Neretti, N., Maioli, T. U., Santos, R. R., Brigidi, P., Franceschi, C., & Faria, A. M. C. (2020). Inflammaging in Endemic Areas for Infectious Diseases. Frontiers in Immunology, 11. https://www.frontiersin.org/articles/10.3389/fimmu.2020.579972

Battram, T., Yousefi, P., Crawford, G., Prince, C., Sheikhali Babaei, M., Sharp, G., Hatcher, C., Vega-Salas, M. J., Khodabakhsh, S., Whitehurst, O., Langdon, R., Mahoney, L., Elliott, H. R., Mancano, G., Lee, M. A., Watkins, S. H., Lay, A. C., Hemani, G., Gaunt, T. R., … Suderman, M. (2022). The EWAS Catalog: A database of epigenome-wide association studies. Wellcome Open Research, 7, 41. 10.12688/wellcomeopenres.17598.2

Beagle, A. J., Tison, G. H., Aschbacher, K., Olgin, J. E., Marcus, G. M., & Pletcher, M. J. (2020). Comparison of the Physical Activity Measured by a Consumer Wearable Activity Tracker and That Measured by Self-Report: Cross-Sectional Analysis of the Health eHeart Study. JMIR MHealth and UHealth, 8(12), e22090. 10.2196/22090

Belsky, D. W., Caspi, A., Corcoran, D. L., Sugden, K., Poulton, R., Arseneault, L., Baccarelli, A., Chamarti, K., Gao, X., Hannon, E., Harrington, H. L., Houts, R., Kothari, M., Kwon, D., Mill, J., Schwartz, J., Vokonas, P., Wang, C., Williams, B. S., & Moffitt, T. E. (2022). DunedinPACE, a DNA methylation biomarker of the pace of aging. ELife, 11. 10.7554/eLife.73420

Brevetti, G., Giugliano, G., Brevetti, L., & Hiatt, W. R. (2010). Inflammation in Peripheral Artery Disease. Circulation, 122(18), 1862–1875. 10.1161/CIRCULATIONAHA.109.918417

Chang, D., Sharma, L., & Dela Cruz, C. S. (2020). Chitotriosidase: A marker and modulator of lung disease. European Respiratory Review, 29(156), 190143. 10.1183/16000617.0143-2019

Christopoulos, P. F., Gjølberg, T. T., Krüger, S., Haraldsen, G., Andersen, J. T., & Sundlisæter, E. (2021). Targeting the Notch Signaling Pathway in Chronic Inflammatory Diseases. Frontiers in Immunology, 12. https://www.frontiersin.org/articles/10.3389/fimmu.2021.668207

Costantini, A., Viola, N., Berretta, A., Galeazzi, R., Matacchione, G., Sabbatinelli, J., Storci, G., De Matteis, S., Butini, L., Rippo, M. R., Procopio, A. D., Caraceni, D., Antonicelli, R., Olivieri, F., & Bonafè, M. (2018). Age-related M1/M2 phenotype changes in circulating monocytes from healthy/unhealthy individuals. Aging, 10(6), 1268–1280. 10.18632/aging.101465

Cui, A., Huang, T., Li, S., Ma, A., Pérez, J. L., Sander, C., Keskin, D. B., Wu, C. J., Fraenkel, E., & Hacohen, N. (2023). Dictionary of immune responses to cytokines at single-cell resolution. Nature. 10.1038/s41586-023-06816-9

De Maeyer, R. P. H., & Chambers, E. S. (2021). The impact of ageing on monocytes and macrophages. Immunology Letters, 230, 1–10. 10.1016/j.imlet.2020.12.003

Edgar, R., Domrachev, M., & Lash, A. E. (2002). Gene Expression Omnibus: NCBI gene expression and hybridization array data repository. Nucleic Acids Research, 30(1), 207–210. 10.1093/nar/30.1.207

Ferrucci, L., & Fabbri, E. (2018). Inflammageing: Chronic inflammation in ageing, cardiovascular disease, and frailty. Nature Reviews. Cardiology, 15(9), 505–522. 10.1038/s41569-018-0064-2

Fortin, J.-P., Fertig, E., & Hansen, K. (2014). shinyMethyl: Interactive quality control of Illumina 450k DNA methylation arrays in R. F1000Research, 3, 175. 10.12688/f1000research.4680.2

Friedman, J., Hastie, T., & Tibshirani, R. (2010). Regularization Paths for Generalized Linear Models via Coordinate Descent. Journal of Statistical Software, 33(1), 1–22.

Furman, D., Campisi, J., Verdin, E., Carrera-Bastos, P., Targ, S., Franceschi, C., Ferrucci, L., Gilroy, D. W., Fasano, A., Miller, G. W., Miller, A. H., Mantovani, A., Weyand, C. M., Barzilai, N., Goronzy, J. J., Rando, T. A., Effros, R. B., Lucia, A., Kleinstreuer, N., & Slavich, G. M. (2019). Chronic inflammation in the etiology of disease across the life span. Nature Medicine, 25(12), 1822–1832. 10.1038/s41591-019-0675-0

Gadd, D. A., Hillary, R. F., McCartney, D. L., Zaghlool, S. B., Stevenson, A. J., Cheng, Y., Fawns-Ritchie, C., Nangle, C., Campbell, A., Flaig, R., Harris, S. E., Walker, R. M., Shi, L., Tucker-Drob, E. M., Gieger, C., Peters, A., Waldenberger, M., Graumann, J., McRae, A. F., … Marioni, R. E. (2022). Epigenetic scores for the circulating proteome as tools for disease prediction. ELife, 11, e71802. 10.7554/eLife.71802

Galkin, F., Kochetov, K., Mamoshina, P., & Zhavoronkov, A. (2021). Adapting Blood DNA Methylation Aging Clocks for Use in Saliva Samples With Cell-type Deconvolution. Frontiers in Aging, 2. https://www.frontiersin.org/articles/10.3389/fragi.2021.697254

Geurts, J. M. W., Janssen, R. G. J. H., van Greevenbroek, M. M. J., van der Kallen, C. J. H., Cantor, R. M., Bu, X., Aouizerat, B. E., Allayee, H., Rotter, J. I., & de Bruin, T. W. A. (2000). Identification of TNFRSF1B as a novel modifier gene in familial combined hyperlipidemia. Human Molecular Genetics, 9(14), 2067–2074. 10.1093/hmg/9.14.2067

Gordon, H. (2001). Detection of alcoholic liver disease. World Journal of Gastroenterology, 7(3), 297–302. 10.3748/wjg.v7.i3.297

Haghani, A., Li, C. Z., Robeck, T. R., Zhang, J., Lu, A. T., Ablaeva, J., Acosta-Rodríguez, V. A., Adams, D. M., Alagaili, A. N., Almunia, J., Aloysius, A., Amor, N. M. S., Ardehali, R., Arneson, A., Baker, C. S., Banks, G., Belov, K., Bennett, N. C., Black, P., … Horvath, S. (2023). DNA methylation networks underlying mammalian traits. Science, 381(6658), eabq5693. 10.1126/science.abq5693

Hannum, G., Guinney, J., Zhao, L., Zhang, L., Hughes, G., Sadda, S., Klotzle, B., Bibikova, M., Fan, J.-B., Gao, Y., Deconde, R., Chen, M., Rajapakse, I., Friend, S., Ideker, T., & Zhang, K. (2013). Genome-wide methylation profiles reveal quantitative views of human aging rates. Molecular Cell, 49(2), 359–367. 10.1016/j.molcel.2012.10.016

Hayashino, Y., Jackson, J. L., Hirata, T., Fukumori, N., Nakamura, F., Fukuhara, S., Tsujii, S., & Ishii, H. (2014). Effects of exercise on C-reactive protein, inflammatory cytokine and adipokine in patients with type 2 diabetes: A meta-analysis of randomized controlled trials. Metabolism - Clinical and Experimental, 63(3), 431–440. 10.1016/j.metabol.2013.08.018

Higgins-Chen, A. T., Thrush, K. L., Wang, Y., Minteer, C. J., Kuo, P.-L., Wang, M., Niimi, P., Sturm, G., Lin, J., Moore, A. Z., Bandinelli, S., Vinkers, C. H., Vermetten, E., Rutten, B. P. F., Geuze, E., Okhuijsen-Pfeifer, C., van der Horst, M. Z., Schreiter, S., Gutwinski, S., … Levine, M. E. (2022). A computational solution for bolstering reliability of epigenetic clocks: Implications for clinical trials and longitudinal tracking. Nature Aging, 2(7), 644–661. 10.1038/s43587-022-00248-2

Hillary, R. F., Trejo-Banos, D., Kousathanas, A., McCartney, D. L., Harris, S. E., Stevenson, A. J., Patxot, M., Ojavee, S. E., Zhang, Q., Liewald, D. C., Ritchie, C. W., Evans, K. L., Tucker-Drob, E. M., Wray, N. R., McRae, A. F., Visscher, P. M., Deary, I. J., Robinson, M. R., & Marioni, R. E. (2020). Multi-method genome- and epigenome-wide studies of inflammatory protein levels in healthy older adults. Genome Medicine, 12(1), 60. 10.1186/s13073-020-00754-1

Horvath, S. (2013). DNA methylation age of human tissues and cell types. Genome Biology, 14(10), R115. 10.1186/gb-2013-14-10-r115

Horvath, S., Oshima, J., Martin, G. M., Lu, A. T., Quach, A., Cohen, H., Felton, S., Matsuyama, M., Lowe, D., Kabacik, S., Wilson, J. G., Reiner, A. P., Maierhofer, A., Flunkert, J., Aviv, A., Hou, L., Baccarelli, A. A., Li, Y., Stewart, J. D., … Raj, K. (2018). Epigenetic clock for skin and blood cells applied to Hutchinson Gilford Progeria Syndrome and ex vivo studies. Aging, 10(7), 1758–1775. 10.18632/aging.101508

Horvath, S., & Raj, K. (2018). DNA methylation-based biomarkers and the epigenetic clock theory of ageing. Nature Reviews Genetics, 19(6), 371–384. 10.1038/s41576-018-0004-3

Jones, M. W., Gnanapandithan, K., Panneerselvam, D., & Ferguson, T. (2023). Chronic Cholecystitis. In StatPearls. StatPearls Publishing.

Kennedy, B. K., Berger, S. L., Brunet, A., Campisi, J., Cuervo, A. M., Epel, E. S., Franceschi, C., Lithgow, G. J., Morimoto, R. I., Pessin, J. E., Rando, T. A., Richardson, A., Schadt, E. E., Wyss-Coray, T., & Sierra, F. (2014). Geroscience: Linking aging to chronic disease. Cell, 159(4), 709–713. 10.1016/j.cell.2014.10.039

Knoppers, T., Beauchamp, E., Dewar, K., Kimmins, S., Bourque, G., Joly, Y., & Dupras, C. (2021). The omics of our lives: Practices and policies of direct-to-consumer epigenetic and microbiomic testing companies. New Genetics and Society, 40(4), 541–569. 10.1080/14636778.2021.1997576

Koelman, L., Pivovarova-Ramich, O., Pfeiffer, A. F. H., Grune, T., & Aleksandrova, K. (2019). Cytokines for evaluation of chronic inflammatory status in ageing research: Reliability and phenotypic characterisation. Immunity & Ageing, 16(1), 11. 10.1186/s12979-019-0151-1

Koestler, D. C., Usset, J., Christensen, B. C., Marsit, C. J., Karagas, M. R., Kelsey, K. T., & Wiencke, J. K. (2017). DNA Methylation-Derived Neutrophil-to-Lymphocyte Ratio: An Epigenetic Tool to Explore Cancer Inflammation and Outcomes. *Cancer Epidemiology, Biomarkers & Prevention : A Publication of the American Association for Cancer Research*, Cosponsored by the American Society of Preventive Oncology, 26(3), 328–338. 10.1158/1055-9965.EPI-16-0461

Kolberg, L., Raudvere, U., Kuzmin, I., Adler, P., Vilo, J., & Peterson, H. (2023). g:Profiler—Interoperable web service for functional enrichment analysis and gene identifier mapping (2023 update). Nucleic Acids Research, 51(W1), W207–W212. 10.1093/nar/gkad347

Koo, T. K., & Li, M. Y. (2016). A Guideline of Selecting and Reporting Intraclass Correlation Coefficients for Reliability Research. Journal of Chiropractic Medicine, 15(2), 155–163. 10.1016/j.jcm.2016.02.012

Kuan, V., Denaxas, S., Patalay, P., Nitsch, D., Mathur, R., Gonzalez-Izquierdo, A., Sofat, R., Partridge, L., Roberts, A., Wong, I. C. K., Hingorani, M., Chaturvedi, N., Hemingway, H., & Hingorani, A. D. (2023). Identifying and visualising multimorbidity and comorbidity patterns in patients in the English National Health Service: A population-based study. The Lancet. Digital Health, 5(1), e16–e27. 10.1016/S2589-7500(22)00187-X

Largman-Chalamish, M., Wasserman, A., Silberman, A., Levinson, T., Ritter, O., Berliner, S., Zeltser, D., Shapira, I., Rogowski, O., & Shenhar-Tsarfaty, S. (2022). Differentiating between bacterial and viral infections by estimated CRP velocity. PLOS ONE, 17(12), e0277401. 10.1371/journal.pone.0277401

Levine, M. E., Lu, A. T., Quach, A., Chen, B. H., Assimes, T. L., Bandinelli, S., Hou, L., Baccarelli, A. A., Stewart, J. D., Li, Y., Whitsel, E. A., Wilson, J. G., Reiner, A. P., Aviv, A., Lohman, K., Liu, Y., Ferrucci, L., & Horvath, S. (2018). An epigenetic biomarker of aging for lifespan and healthspan. Aging, 10(4), 573–591. 10.18632/aging.101414

Li, M., Zou, D., Li, Z., Gao, R., Sang, J., Zhang, Y., Li, R., Xia, L., Zhang, T., Niu, G., Bao, Y., & Zhang, Z. (2019). EWAS Atlas: A curated knowledgebase of epigenome-wide association studies. Nucleic Acids Research, 47(D1), D983–D988. 10.1093/nar/gky1027

Ligthart, S., Marzi, C., Aslibekyan, S., Mendelson, M. M., Conneely, K. N., Tanaka, T., Colicino, E., Waite, L. L., Joehanes, R., Guan, W., Brody, J. A., Elks, C., Marioni, R., Jhun, M. A., Agha, G., Bressler, J., Ward-Caviness, C. K., Chen, B. H., Huan, T., … Dehghan, A. (2016). DNA methylation signatures of chronic low-grade inflammation are associated with complex diseases. Genome Biology, 17(1). 10.1186/s13059-016-1119-5

López-Otín, C., Blasco, M. A., Partridge, L., Serrano, M., & Kroemer, G. (2023). Hallmarks of aging: An expanding universe. Cell, 186(2), 243–278. 10.1016/j.cell.2022.11.001

Lu, A. T., Fei, Z., Haghani, A., Robeck, T. R., Zoller, J. A., Li, C. Z., Lowe, R., Yan, Q., Zhang, J., Vu, H., Ablaeva, J., Acosta-Rodriguez, V. A., Adams, D. M., Almunia, J., Aloysius, A., Ardehali, R., Arneson, A., Baker, C. S., Banks, G., … Horvath, S. (2023). Universal DNA methylation age across mammalian tissues. Nature Aging, 3(9), 1144–1166. 10.1038/s43587-023-00462-6

Lu, A. T., Quach, A., Wilson, J. G., Reiner, A. P., Aviv, A., Raj, K., Hou, L., Baccarelli, A. A., Li, Y., Stewart, J. D., Whitsel, E. A., Assimes, T. L., Ferrucci, L., & Horvath, S. (2019). DNA methylation GrimAge strongly predicts lifespan and healthspan. Aging, 11(2), 303–327. 10.18632/aging.101684

Mack, I., Hector, A., Ballbach, M., Kohlhäufl, J., Fuchs, K. J., Weber, A., Mall, M. A., & Hartl, D. (2015). The role of chitin, chitinases, and chitinase-like proteins in pediatric lung diseases. Molecular and Cellular Pediatrics, 2(1), 3. 10.1186/s40348-015-0014-6

Makgoeng, S. B., Bolanos, R. S., Jeon, C. Y., Weiss, R. E., Arah, O. A., Breen, E. C., Martínez-Maza, O., & Hussain, S. K. (2018). Markers of Immune Activation and Inflammation, and Non-Hodgkin Lymphoma: A Meta-Analysis of Prospective Studies. JNCI Cancer Spectrum, 2(4), pky082. 10.1093/jncics/pky082

Margiotti, K., Monaco, F., Fabiani, M., Mesoraca, A., & Giorlandino, C. (2023). Epigenetic Clocks: In Aging-Related and Complex Diseases. Cytogenetic and Genome Research, 1–10. 10.1159/000534561

Márquez, E. J., Chung, C., Marches, R., Rossi, R. J., Nehar-Belaid, D., Eroglu, A., Mellert, D. J., Kuchel, G. A., Banchereau, J., & Ucar, D. (2020). Sexual-dimorphism in human immune system aging. Nature Communications, 11(1), 751. 10.1038/s41467-020-14396-9

Martínez de Toda, I., González-Sánchez, M., Díaz-Del Cerro, E., Valera, G., Carracedo, J., & Guerra-Pérez, N. (2023). Sex differences in markers of oxidation and inflammation. Implications for ageing. Mechanisms of Ageing and Development, 211, 111797. 10.1016/j.mad.2023.111797

Martin-Herranz, D. E. (2019). On the epigenetic ageing clock in humans [PhD thesis, University of Cambridge]. https://api.semanticscholar.org/CorpusID:209584461

Martin-Herranz, D. E., Aref-Eshghi, E., Bonder, M. J., Stubbs, T. M., Choufani, S., Weksberg, R., Stegle, O., Sadikovic, B., Reik, W., & Thornton, J. M. (2019). Screening for genes that accelerate the epigenetic aging clock in humans reveals a role for the H3K36 methyltransferase NSD1. Genome Biology, 20(1), 146. 10.1186/s13059-019-1753-9

Min, J. L., Hemani, G., Davey Smith, G., Relton, C., & Suderman, M. (2018). Meffil: Efficient normalization and analysis of very large DNA methylation datasets. Bioinformatics (Oxford, England), 34(23), 3983–3989. 10.1093/bioinformatics/bty476

Moqri, M., Herzog, C., Poganik, J. R., Justice, J., Belsky, D. W., Higgins-Chen, A., Moskalev, A., Fuellen, G., Cohen, A. A., Bautmans, I., Widschwendter, M., Ding, J., Fleming, A., Mannick, J., Han, J.-D. J., Zhavoronkov, A., Barzilai, N., Kaeberlein, M., Cummings, S., … Gladyshev, V. N. (2023). Biomarkers of aging for the identification and evaluation of longevity interventions. Cell, 186(18), 3758–3775. 10.1016/j.cell.2023.08.003

Nilsonne, G., Lekander, M., Åkerstedt, T., Axelsson, J., & Ingre, M. (2016). Diurnal Variation of Circulating Interleukin-6 in Humans: A Meta-Analysis. PloS One, 11(11), e0165799. 10.1371/journal.pone.0165799

Nishitani, S., Isozaki, M., Yao, A., Higashino, Y., Yamauchi, T., Kidoguchi, M., Kawajiri, S., Tsunetoshi, K., Neish, H., Imoto, H., Arishima, H., Kodera, T., Fujisawa, T. X., Nomura, S., Kikuta, K., Shinozaki, G., & Tomoda, A. (2023). Cross-tissue correlations of genome-wide DNA methylation in Japanese live human brain and blood, saliva, and buccal epithelial tissues. Translational Psychiatry, 13(1), 72. 10.1038/s41398-023-02370-0

Oh, H. S.-H., Rutledge, J., Nachun, D., Pálovics, R., Abiose, O., Moran-Losada, P., Channappa, D., Urey, D. Y., Kim, K., Sung, Y. J., Wang, L., Timsina, J., Western, D., Liu, M., Kohlfeld, P., Budde, J., Wilson, E. N., Guen, Y., Maurer, T. M., … Wyss-Coray, T. (2023). Organ aging signatures in the plasma proteome track health and disease. Nature, 624(7990), 164–172. 10.1038/s41586-023-06802-1

Oudijk, E. D., Lammers, J. J., & Koenderman, L. (2003). Systemic inflammation in chronic obstructive pulmonary disease. European Respiratory Journal, *22*(46 suppl), 5s. 10.1183/09031936.03.00004603a

Parkinson, H., Kapushesky, M., Shojatalab, M., Abeygunawardena, N., Coulson, R., Farne, A., Holloway, E., Kolesnykov, N., Lilja, P., Lukk, M., Mani, R., Rayner, T., Sharma, A., William, E., Sarkans, U., & Brazma, A. (2007). ArrayExpress—A public database of microarray experiments and gene expression profiles. Nucleic Acids Research, 35(Database issue), D747-750. 10.1093/nar/gkl995

Petagna, L., Antonelli, A., Ganini, C., Bellato, V., Campanelli, M., Divizia, A., Efrati, C., Franceschilli, M., Guida, A. M., Ingallinella, S., Montagnese, F., Sensi, B., Siragusa, L., & Sica, G. S. (2020). Pathophysiology of Crohn’s disease inflammation and recurrence. Biology Direct, 15(1), 23. 10.1186/s13062-020-00280-5

Picard, E., Armstrong, S., Andrew, M. K., Haynes, L., Loeb, M., Pawelec, G., Kuchel, G. A., McElhaney, J. E., & Verschoor, C. P. (2022). Markers of systemic inflammation are positively associated with influenza vaccine antibody responses with a possible role for ILT2(+)CD57(+) NK-cells. Immunity & Ageing, 19(1), 26. 10.1186/s12979-022-00284-x

Pidsley, R., Y Wong, C. C., Volta, M., Lunnon, K., Mill, J., & Schalkwyk, L. C. (2013). A data-driven approach to preprocessing Illumina 450K methylation array data. BMC Genomics, 14(1), 293. 10.1186/1471-2164-14-293

Quach, A., Levine, M. E., Tanaka, T., Lu, A. T., Chen, B. H., Ferrucci, L., Ritz, B., Bandinelli, S., Neuhouser, M. L., Beasley, J. M., Snetselaar, L., Wallace, R. B., Tsao, P. S., Absher, D., Assimes, T. L., Stewart, J. D., Li, Y., Hou, L., Baccarelli, A. A., … Horvath, S. (2017). Epigenetic clock analysis of diet, exercise, education, and lifestyle factors. Aging, 9(2), 419–446. 10.18632/aging.101168

R Core Team. (2023). *R: A Language and Environment for Statistical Computing*. R Foundation for Statistical Computing. https://www.R-project.org/

Rodriguez-Meira, A., Norfo, R., Wen, S., Chédeville, A. L., Rahman, H., O’Sullivan, J., Wang, G., Louka, E., Kretzschmar, W. W., Paterson, A., Brierley, C., Martin, J.-E., Demeule, C., Bashton, M., Sousos, N., Moralli, D., Subha Meem, L., Carrelha, J., Wu, B., … Mead, A. J. (2023). Single-cell multi-omics identifies chronic inflammation as a driver of TP53-mutant leukemic evolution. Nature Genetics, 55(9), 1531–1541. 10.1038/s41588-023-01480-1

Saffari, A., Silver, M. J., Zavattari, P., Moi, L., Columbano, A., Meaburn, E. L., & Dudbridge, F. (2018). Estimation of a significance threshold for epigenome-wide association studies. Genetic Epidemiology, 42(1), 20–33. 10.1002/gepi.22086

Saule, P., Trauet, J., Dutriez, V., Lekeux, V., Dessaint, J.-P., & Labalette, M. (2006). Accumulation of memory T cells from childhood to old age: Central and effector memory cells in CD4+ versus effector memory and terminally differentiated memory cells in CD8+ compartment. Mechanisms of Ageing and Development, 127(3), 274–281. 10.1016/j.mad.2005.11.001

Sayed, N., Huang, Y., Nguyen, K., Krejciova-Rajaniemi, Z., Grawe, A. P., Gao, T., Tibshirani, R., Hastie, T., Alpert, A., Cui, L., Kuznetsova, T., Rosenberg-Hasson, Y., Ostan, R., Monti, D., Lehallier, B., Shen-Orr, S. S., Maecker, H. T., Dekker, C. L., Wyss-Coray, T., … Furman, D. (2021). An inflammatory aging clock (iAge) based on deep learning tracks multimorbidity, immunosenescence, frailty and cardiovascular aging. Nature Aging, 1(7), 598–615. 10.1038/s43587-021-00082-y

Schraufstatter, I. U., Zhao, M., Khaldoyanidi, S. K., & Discipio, R. G. (2012). The chemokine CCL18 causes maturation of cultured monocytes to macrophages in the M2 spectrum. Immunology, 135(4), 287–298. 10.1111/j.1365-2567.2011.03541.x

Scott, A. J., Ellison, M., & Sinclair, D. A. (2021). The economic value of targeting aging. Nature Aging, 1(7), 616–623. 10.1038/s43587-021-00080-0

Siemons, L., Ten Klooster, P. M., Vonkeman, H. E., van Riel, P. L. C. M., Glas, C. A. W., & van de Laar, M. A. F. J. (2014). How age and sex affect the erythrocyte sedimentation rate and C-reactive protein in early rheumatoid arthritis. BMC Musculoskeletal Disorders, 15, 368. 10.1186/1471-2474-15-368

Sierra, F. (2016). The Emergence of Geroscience as an Interdisciplinary Approach to the Enhancement of Health Span and Life Span. Cold Spring Harbor Perspectives in Medicine, 6(4), a025163. 10.1101/cshperspect.a025163

Smith, B. H., Campbell, A., Linksted, P., Fitzpatrick, B., Jackson, C., Kerr, S. M., Deary, I. J., MacIntyre, D. J., Campbell, H., McGilchrist, M., Hocking, L. J., Wisely, L., Ford, I., Lindsay, R. S., Morton, R., Palmer, C. N. A., Dominiczak, A. F., Porteous, D. J., & Morris, A. D. (2013). Cohort Profile: Generation Scotland: Scottish Family Health Study (GS:SFHS). The study, its participants and their potential for genetic research on health and illness. International Journal of Epidemiology, 42(3), 689–700. 10.1093/ije/dys084

Štambuk, J., Nakić, N., Vučković, F., Pučić-Baković, M., Razdorov, G., Trbojević-Akmačić, I., Novokmet, M., Keser, T., Vilaj, M., Štambuk, T., Gudelj, I., Šimurina, M., Song, M., Wang, H., Salihović, M. P., Campbell, H., Rudan, I., Kolčić, I., Eller, L. A., … Lauc, G. (2020). Global variability of the human IgG glycome. Aging, 12(15), 15222–15259. 10.18632/aging.103884

Stevenson, A. J., Gadd, D. A., Hillary, R. F., McCartney, D. L., Campbell, A., Walker, R. M., Evans, K. L., Harris, S. E., Spires-Jones, T. L., McRae, A. F., Visscher, P. M., McIntosh, A. M., Deary, I. J., & Marioni, R. E. (2021). Creating and Validating a DNA Methylation-Based Proxy for Interleukin-6. Journals of Gerontology - Series A Biological Sciences and Medical Sciences, 76(12), 2284–2292. 10.1093/gerona/glab046

Stevenson, A. J., McCartney, D. L., Hillary, R. F., Campbell, A., Morris, S. W., Bermingham, M. L., Walker, R. M., Evans, K. L., Boutin, T. S., Hayward, C., McRae, A. F., McColl, B. W., Spires-Jones, T. L., McIntosh, A. M., Deary, I. J., & Marioni, R. E. (2020). Characterisation of an inflammation-related epigenetic score and its association with cognitive ability. Clinical Epigenetics, 12(1). 10.1186/s13148-020-00903-8

System COVID-19 Codes – Available March 2020 Coding – Primary Care Informatics. (2020, April 9). SCIMP. https://www.scimp.scot.nhs.uk/archives/2733

Szabo, Y. Z., & Slavish, D. C. (2021). Measuring salivary markers of inflammation in health research: A review of methodological considerations and best practices. Psychoneuroendocrinology, 124, 105069. 10.1016/j.psyneuen.2020.105069

Tang, Y., Liang, P., Chen, J., Fu, S., Liu, B., Feng, M., Lin, B., Lee, B., Xu, A., & Lan, H. Y. (2018). The baseline levels and risk factors for high-sensitive C-reactive protein in Chinese healthy population. Immunity & Ageing, 15(1), 21. 10.1186/s12979-018-0126-7

Teschendorff, A. E., Marabita, F., Lechner, M., Bartlett, T., Tegner, J., Gomez-Cabrero, D., & Beck, S. (2013). A beta-mixture quantile normalization method for correcting probe design bias in Illumina Infinium 450 k DNA methylation data. *Bioinformatics (Oxford*, England), 29(2), 189–196. 10.1093/bioinformatics/bts680

Tomiyama, H., Matsuda, T., & Takiguchi, M. (2002). Differentiation of Human CD8+ T Cells from a Memory to Memory/Effector Phenotype1. The Journal of Immunology, 168(11), 5538–5550. 10.4049/jimmunol.168.11.5538

Triche, T. J., Jr, Weisenberger, D. J., Van Den Berg, D., Laird, P. W., & Siegmund, K. D. (2013). Low-level processing of Illumina Infinium DNA Methylation BeadArrays. Nucleic Acids Research, 41(7), e90–e90. 10.1093/nar/gkt090

van Dongen, J., Nivard, M. G., Willemsen, G., Hottenga, J.-J., Helmer, Q., Dolan, C. V., Ehli, E. A., Davies, G. E., van Iterson, M., Breeze, C. E., Beck, S., BIOS Consortium, Suchiman, H. E., Jansen, R., van Meurs, J. B., Heijmans, B. T., Slagboom, P. E., & Boomsma, D. I. (2016). Genetic and environmental influences interact with age and sex in shaping the human methylome. Nature Communications, 7, 11115. 10.1038/ncomms11115

Vazirinejad, R., Ahmadi, Z., Kazemi Arababadi, M., Hassanshahi, G., & Kennedy, D. (2014). The Biological Functions, Structure and Sources of CXCL10 and Its Outstanding Part in the Pathophysiology of Multiple Sclerosis. Neuroimmunomodulation, 21(6), 322–330. 10.1159/000357780

Wang, Y.-C., He, F., Feng, F., Liu, X.-W., Dong, G.-Y., Qin, H.-Y., Hu, X.-B., Zheng, M.-H., Liang, L., Feng, L., Liang, Y.-M., & Han, H. (2010). Notch Signaling Determines the M1 versus M2 Polarization of Macrophages in Antitumor Immune Responses. Cancer Research, 70(12), 4840–4849. 10.1158/0008-5472.CAN-10-0269

Xu, Z., & Taylor, J. A. (2021). Reliability of DNA methylation measures using Illumina methylation BeadChip. Epigenetics, 16(5), 495–502. 10.1080/15592294.2020.1805692

Zheng, S. C., Webster, A. P., Dong, D., Feber, A., Graham, D. G., Sullivan, R., Jevons, S., Lovat, L. B., Beck, S., Widschwendter, M., & Teschendorff, A. E. (2018). A novel cell-type deconvolution algorithm reveals substantial contamination by immune cells in saliva, buccal and cervix. Epigenomics, 10(7), 925–940. 10.2217/epi-2018-0037

Zhou, W., Triche, T. J., Jr, Laird, P. W., & Shen, H. (2018). SeSAMe: Reducing artifactual detection of DNA methylation by Infinium BeadChips in genomic deletions. Nucleic Acids Research, 46(20), e123–e123. 10.1093/nar/gky691

